# FUS and TAF15 safeguard the critical functions of the ribonucleoprotein network formed by EWSR1 and newly synthesized RNA

**DOI:** 10.64898/2026.03.24.713985

**Authors:** Soumya Sundara Rajan, Imran Khan, Tamara L. Jones, Tayvia Brownmiller, Vernon J. Ebegboni, Langston Lim, Andy Tran, Michael J. Kruhlak, Natasha J. Caplen

## Abstract

The FET family of RNA-binding proteins, FUS, EWSR1, and TAF15, contribute to transcriptional regulation and RNA maturation, but their core functions remain unclear. Chromosomal rearrangements involving FUS, EWSR1, or TAF15 drive multiple cancers, and mutations in the genes encoding the FET proteins are associated with neurodegenerative disease. Here, using nanoscale imaging, we show that endogenous EWSR1 and newly synthesized RNA exhibit a network-like organization with EWSR1 foci forming the nodes of this ribonucleoprotein network. Acute depletion of EWSR1 causes a rapid but transient reduction in nascent RNA levels and cellular metabolic activity without affecting active transcription. Notably, loss of EWSR1 induces a compensatory mechanism involving the reorganization of FUS and TAF15 to closely resemble that of EWSR1, including enhanced clustering with newly synthesized RNA. Together, our findings reveal functional redundancy within the FET protein family that is critical for the homeostatic regulation of nascent RNA levels.

**In brief:** Sundara Rajan et al. show that endogenous EWSR1 and nascent RNA form a ribonucleoprotein network. EWSR1 depletion transiently reduces nascent RNA and metabolic activity without impairment of transcriptional elongation. Loss of EWSR1 induces compensatory reorganization of FUS and TAF15, revealing a protein family mechanism required for the homeostatic regulation of nascent RNA levels.

**Highlights:**

- EWSR1 and nascent RNA form a ribonucleoprotein network
- EWSR1 loss transiently reduces nascent RNA and metabolic activity
- FUS and TAF15 undergo compensatory nuclear reorganization upon EWSR1 loss
- FUS and TAF15 functionally replace EWSR1

**GRAPHICAL ABSTRACT:** 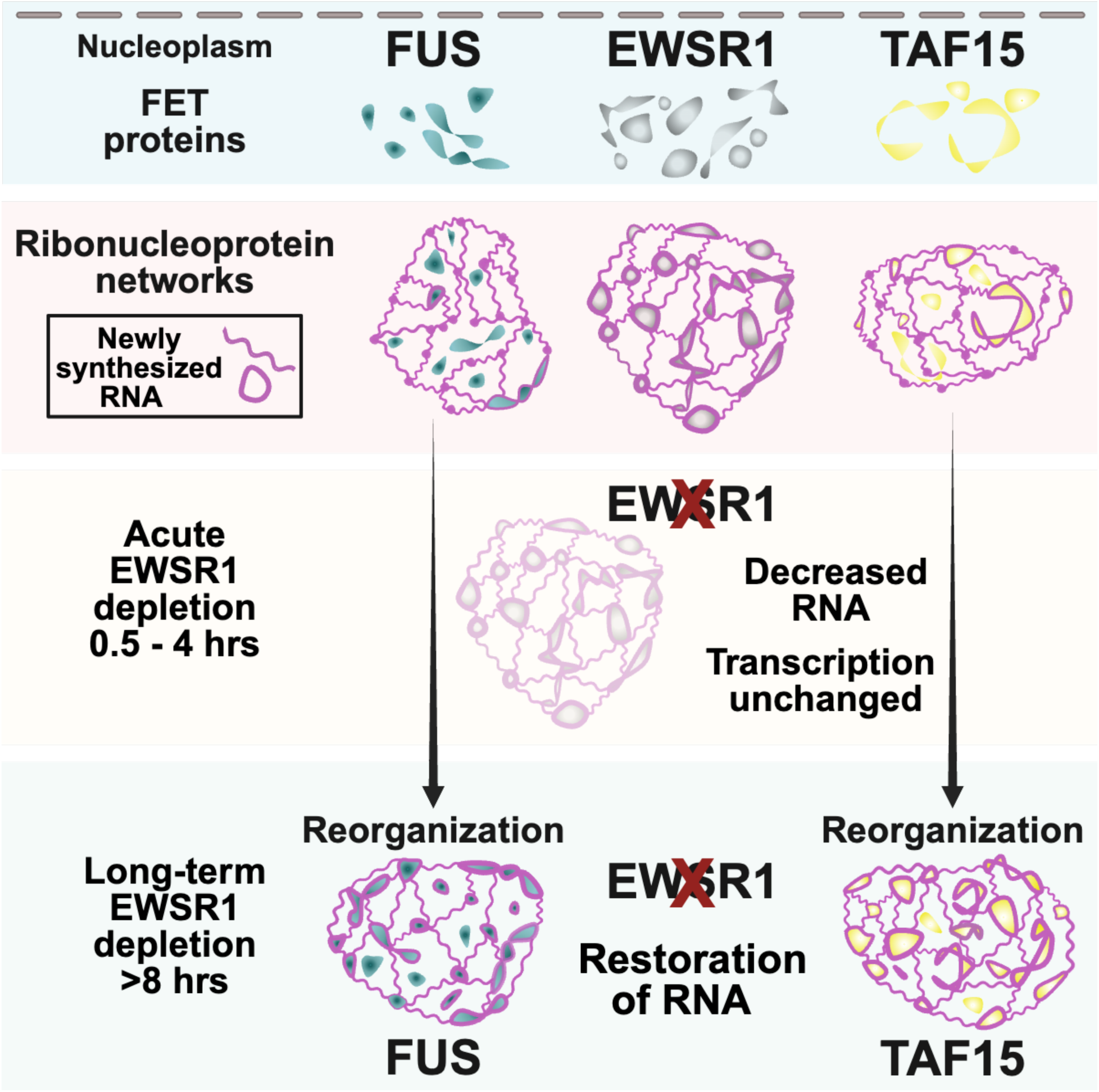

## INTRODUCTION

FUS (Fused in Sarcoma), EWSR1 (EWS RNA Binding Protein 1), and TAF15 (TATA-Binding Protein-Associated Factor 2N) comprise the heterogeneous ribonucleoprotein (hnRNP) P family.^1–4^ Collectively referred to as the FET proteins, these RNA-binding proteins (RBPs) are highly homologous and share a conserved domain architecture, including an N-terminal low complexity domain (LCD) enriched in serine, tyrosine, glycine, and glutamine residues.^5^ *In vitro* studies have shown that such LCDs can engage in the multivalent interactions that promote biomolecular condensate formation.^6^ The FET proteins also contain multiple arginine and glycine rich domains that flank an RNA recognition motif (RRM) and a zinc-finger binding domain, along with a C-terminal non-classical pY nuclear localization signal. These shared structural features suggest overlapping molecular functions, yet there is evidence that the individual FET proteins have distinct biological roles.

Elucidating the biological functions of the FET proteins is of particular importance given their links to human disease. Several neurodegenerative disorders including frontotemporal dementia and amyotrophic lateral sclerosis (ALS) involve mutations in genes encoding the FET proteins, most prominently FUS and TAF15.^7,8^ In affected individuals, disease-associated mutations result in protein mislocalization and cytoplasmic accumulation, which correlate with the formation of pathological plaques and fibrillar aggregates.^9,10^ These pathogenic phenotypes reflect a toxic gain-of-function mechanism as Fus^-\+^ mice do not develop ALS.^11^ The FET proteins also play important roles in several cancers, particularly sarcomas affecting children and young adults, in which a chromosomal translocation involving the regulatory elements and sequences encoding the N-terminus of a FET protein become fused to sequences coding for the C-terminal functional domain of a partner protein. These genomic rearrangements result in the expression of a fusion oncoprotein that promotes malignant transformation and tumor cell proliferation. Examples of FET-fusion oncoproteins include TAF15::ZNF384 in acute lymphoblastic leukemia, FUS/EWSR1::DDIT3 in myxoid liposarcoma, EWSR1/FUS/TAF15::NR4A3 in extra skeletal myxoid chondrosarcoma, EWSR1::WT1 in desmoplastic small round cell tumors, and EWSR1::FLI1/ERG/FEV and FUS::FEV in Ewing sarcoma (EWS).^12–17^

Biochemical and functional studies of the FET proteins have converged on two potentially interconnected properties: their association with RNA and their proposed functions in regulating transcription through interactions with RNA polymerase II (RNA pol II). Although the FET protein RRMs do not bind a strict consensus RNA sequence, they may preferentially associate with RNA motifs enriched for specific residues that influence RNA structure.^18,19^ In parallel with their RNA-binding capacity, multiple biochemical studies have demonstrated direct interactions between FET proteins and the C-terminal domain (CTD) of POLR2A, the largest subunit of RNA pol II.^3,20–23^ Post-translational modifications (PTMs) of the YSPTSPS heptapeptide repeats within the POLR2A CTD govern the transcriptional dynamics of RNA pol II. Specifically, phosphorylation of serine 5 (pS5) by CDK7 regulates transcription initiation, whereas phosphorylation of serine 2 (pS2) by CDK9 marks the elongation phase of transcription.^24^ Biochemical studies have proposed a model in which the N-terminus of the FET proteins interact with the CTD of POLR2A, which also contains an LCD, to promote the formation of high concentration hubs that can recruit RNA pol II and facilitate transcriptional regulation.^20,21,23^ Interestingly, analyses of EWSR1::FLI1 have suggested that the fusion oncoprotein can disrupt EWSR1’s regulation of the phosphorylation of the POL2RA CTD.^25^ However, the precise contributions of individual family members and the relevance of these interactions in the context of the full-length endogenous proteins expressed at physiological concentrations remains incompletely understood.

To begin to address this gap in knowledge, we previously characterized the nuclear organization of EWSR1 using two EWS cell lines engineered to express endogenous EWSR1 fused to a fluorescent reporter.^26^ Using these cell lines, we identified EWSR1 as existing in two distinct states: a lower intensity diffused state and a higher-intensity focal state. We demonstrated that in the nucleus there is exclusion of EWSR1 from condensed regions of DNA, that the transcriptional coactivators BRD4 and MED1 flank focal EWSR1 signals, and that these EWSR1 foci showed substantial overlap with phosphorylated RNA-pol II (pS5 and pS2-RNA pol II). Furthermore, we observed nascent RNA in proximity of EWSR1 in both diffused and focal states.

Here, we confirm and extend these observations using non-EWS and EWS cells, nanoscale resolution STED microscopy, and a comprehensive analysis of the consequences of EWSR1 depletion. Our results establish that EWSR1 is an integral component of a ribonucleoprotein (RNP) network. Acute depletion of EWSR1 leads to a rapid reduction in nascent RNA and cellular metabolic activity. Critically, analysis of pS2-RNA pol II indicates that there is preservation of global transcriptional output following EWSR1 loss. Instead, accompanying depletion of EWSR1 is a marked change in the nuclear organization of its homologs FUS and TAF15 such that their architecture recapitulates that of EWSR1. Collectively, our findings support a model in which EWSR1 forms a scaffold for newly synthesized RNA, with FUS and TAF15 acting as safeguards to preserve this fundamental function in the absence of EWSR1.

## RESULTS

### The spatial organization of endogenous EWSR1 is non-random and conserved across cell types

To investigate the spatial organization of endogenous EWSR1, we analyzed six cell lines representing distinct biological contexts: an epithelial cell line (HEK-293T), a fibrosarcoma line lacking a FET gene rearrangement (HT-1080), and four EWS cell lines expressing wild-type EWSR1 and either EWSR1::FLI1 (A673, TC-32, SK-N-MC) or EWSR1::ERG (TC-106) (**Figure 1A**). Immunoblotting confirmed expression of full-length EWSR1 protein (∼68.5-69 kDa) and multiple lower molecular weight isoforms in all cell lines, as well as the expected EWSR1-fusion oncoprotein in EWS cells (**Figures 1B** and **S1A**).

**Figure 1:**
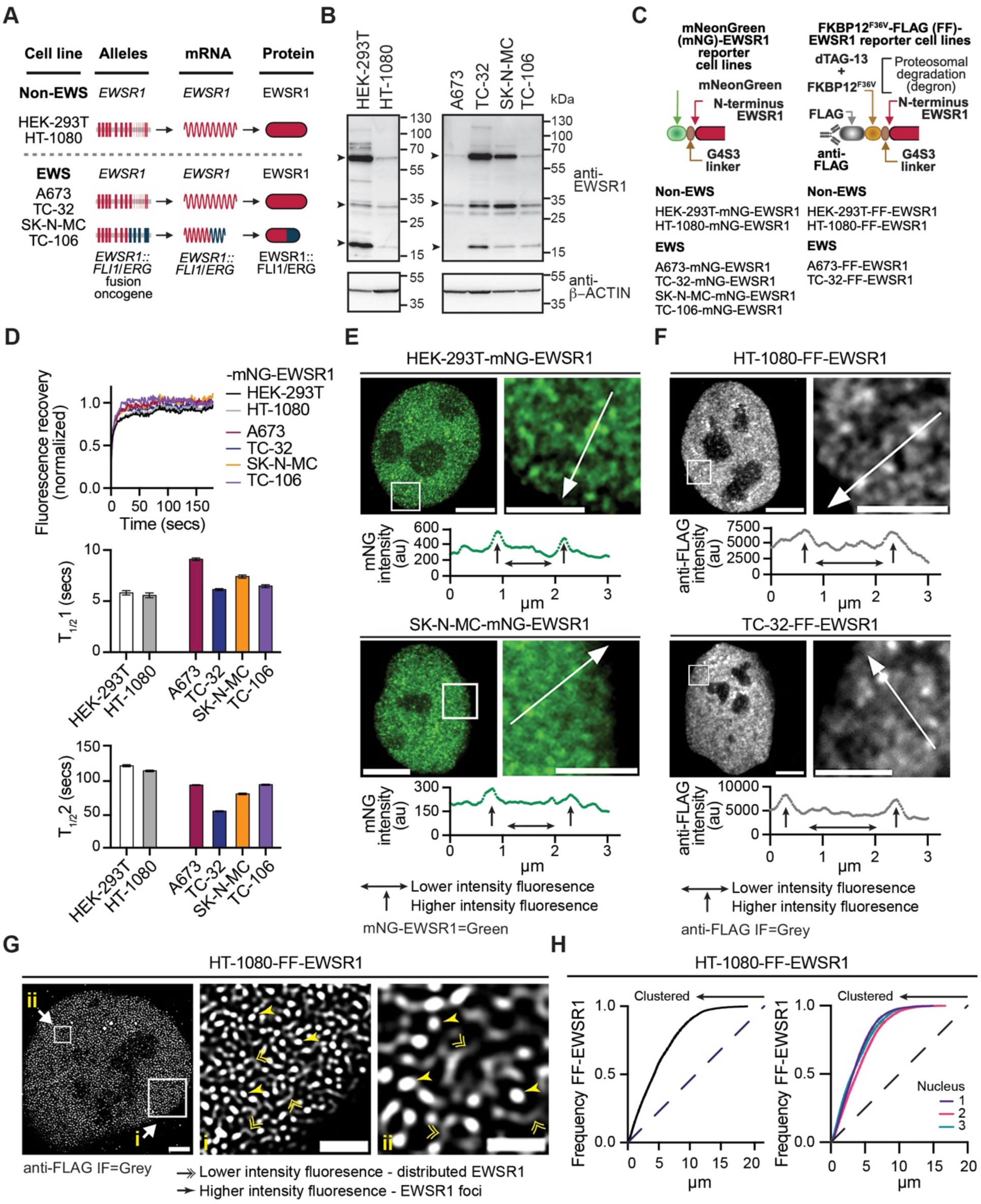
The organization of endogenous EWSR1 is non-random and conserved across cell types. (**A**) Schematic representations of the *EWSR1* alleles and corresponding expressed protein products in the cell lines used in this study. (**B**) Immunoblot analysis of whole-cell lysates from the indicated cell lines probed with antibodies against the indicated proteins. Arrowheads denote EWSR1 isoforms of ∼68.5–69 kDa (UniProt Q01844-1, -3, and -5), an ∼32 kDa isoform (UniProt H7BY36), and ∼16–19 kDa isoforms (UniProt B0QYJ6, B0QYJ4, and B0QYJ3). (**C**) Schematic illustrating N-terminal modifications of endogenous EWSR1 and the nomenclature of the modified EWSR1 reporter cell lines used in this study. Generation and validation of mNeonGreen–EWSR1 (mNG–EWSR1) and FLAG–FKBP12^F36V^–EWSR1 (FF–EWSR1) reporter cell lines are described in **Figures S1** and **S2**, respectively. (**D**) Mean normalized fluorescence recovery after photobleaching (FRAP) recovery curves for mNG–EWSR1 reporter cell lines. Summary kinetics (seconds, secs) for the fast (T_1/2_1) and slow (T_1/2_2) recovering fractions are shown as mean ± SEM (10 bleached regions per cell line). (**E**) SoRa super-resolution images of HEK-293T–mNG–EWSR1 and SK-N-MC–mNG–EWSR1 cells showing nuclei (left) and expanded regions (right). mNG–EWSR1 fluorescence is pseudocolored green. Line-scan intensity plots correspond to the white lines shown in the expanded regions (arrow direction indicates x-axis orientation). Horizontal arrowed lines indicate lower-intensity signal, and vertical arrowed lines indicate higher-intensity signal. Scale bars, nucleus 4 µm, expanded region, 2 µm. (**F**) SoRa super-resolution images of HT-1080–FF–EWSR1 and TC-32–FF–EWSR1 reporter cell lines showing nuclei (left) and expanded regions (right). Anti-FLAG immunofluorescence (IF) is pseudocolored grey. Line-scan intensity plots correspond to the white lines shown in the expanded regions. Scale bars, nucleus 4 µm, expanded region 2 µm. (**G**) STED microscopy anti-FLAG IF (grey) image of an HT-1080–FF–EWSR1 nucleus (left) with two expanded regions (i and ii). Double arrowheads indicate lower-intensity distributed FF–EWSR1 signal and single arrowheads indicate high-intensity FF–EWSR1 foci. Scale bars, nucleus 2 µm, expanded regions i, 1 µm, ii, 0.5 µm. (**H**) Spatial cluster analysis of FF–EWSR1 pattern distribution for a representative HT-1080–FF–EWSR1 nucleus (left) and three additional nuclei (right). The dashed line represents a random point pattern distribution; arrowed solid lines indicate deviation from a random point pattern distribution. Images in **(E–G)** are representative of >20 nuclei per reporter cell line. Schematics in **(A)** and **(C)** were generated using BioRender.

To visualize endogenous EWSR1 without over-expression, we used CRISPR-Cas9-mediated genome editing to insert a DNA cassette at the 5’ end of *EWSR1*-exon 1 encoding either the fluorescence reporter protein mNeonGreen (mNG) or a FLAG-FKBP12^F36V^ (FF) tag (**Figure S1B**). The mNG reporter enabled direct imaging of endogenous EWSR1 at a spatial resolution of ∼120 nm, whereas the FF tag enabled the nanoscale immunofluorescence (IF) imaging (<50 nm spatial resolution) and acute, ligand-inducible proteasomal degradation using the dTAG-13 degron system.^27,28^ Following single-cell cloning, we established two complementary panels of N-terminally tagged EWSR1 reporter cell lines, a complete set of six mNG-EWSR1 expressing cell lines and a subset of four FF-EWSR1 expressing cell lines (**Figure 1C**).

PCR analysis of genomic DNA and sequencing confirmed the correct introduction of the mNG reporter into the unrearranged *EWSR1* locus. We determined monoallelic modification in the HEK-293T and EWS cells, while HT-1080 cells exhibited biallelic modification (**Figure S1C**). All mNG-EWSR1 lines expressed mNG (**Figure S1D**). Importantly, siRNA-mediated depletion of EWSR1, but not EWSR1-fusion transcripts, selectively reduced mNG fluorescence, confirming specific tagging of endogenous EWSR1 (**Figures S1E** and **S1F**, and^26^). Confocal and super-resolution confocal imaging demonstrated restriction of EWSR1 to the nucleus in both non-EWS and EWS cells (**Figures S1G** and **S1H**), consistent with prior reports of endogenous EWSR1’s localization reported as part of the RBP image database.^29^

To assess the dynamic behavior of endogenous EWSR1 and infer its interactions with other macromolecules, we performed fluorescence recovery after photobleaching (FRAP) in the mNG-EWSR1 reporter cell lines. Previous analysis in A673- and TC-32-mNG-EWSR1 reporter cell lines identified two kinetically distinct EWSR1 fractions, a rapidly recovering fraction (T_1/2_ 1: <10 seconds (secs)) and a slower recovering fraction (T_1/2_ 2: >50 secs). Extending this analysis to non-EWS cells and additional EWS cell lines, normalized recovery curves and kinetic fitting confirmed that EWSR1 exists in faster (5.7 – 9.23 secs) and slower (55.75 – 122.6 secs) recovering fractions (**Figure 1D**). The comparable kinetic behavior across the non-EWS and EWS cell lines indicates that EWSR1 engages in dynamic and more stable nuclear interactions independent of the expression of an EWSR1-fusion oncoprotein.

We previously proposed that the presence of two kinetically distinct EWSR1 fractions reflects differential interaction with nucleic acid and other nuclear proteins. In agreement with this model, imaging of A673- and TC-32-mNG-EWSR1 reporter cells revealed two spatially distinct fluorescence states: a diffuse low intensity signal throughout the nucleoplasm and numerous focal regions of high intensity signal. Here, we confirmed and extended these observations to two non-EWS and additional EWS cell lines with the representative line plots of mNG-EWSR1 fluorescence intensity over distance showing the low level nucleoplasmic signal above background (horizontal arrows), interspersed by discrete regions of elevated fluorescence intensity (vertical arrows) (**Figures 1E** and **S1I**). These observations are consistent with the coexistence of diffuse and locally enriched EWSR1.

To independently validate the distinctive organization of endogenous EWSR1 observed using the mNG-EWSR1 reporter cell lines, we next examined the FF-EWSR1 reporter cell lines. We confirmed correct in-frame integration of the FF cassette at *EWSR1* by PCR analysis of genomic DNA and expressed transcripts. HEK-293T and HT-1080 clones exhibited biallelic modification, while in the EWS cell lines, PCR analysis detected modification of only the unrearranged *EWSR1* (**Figures S2A** and **S2B**). Anti-FLAG immunoblotting detected modification of multiple EWSR1 isoforms (**Figure S2C**) and IF analysis confirmed exclusive nuclear localization of FF-EWSR1 (**Figures S2D** and **S2E**). Notably, IF imaging of FF-EWSR1 recapitulated the distinct states of EWSR1 observed in mNG EWSR1 cells as illustrated by the representative images and line plots shown in **Figure 1F** and **Figure S2F**.

To further resolve EWSR1 organization at nanoscale resolution limits, we performed anti-FLAG IF coupled with STED microscopy achieving sub-50 nm spatial resolution. STED images revealed low-intensity signal (double arrowheads) interspersed with discrete areas of high intensity signal (single arrowheads) (**Figures 1G** and **S2G**). Hereafter, we refer to these low and high intensity signals as distributed EWSR1 and EWSR1 foci, respectively.

To quantify the spatial organization of endogenous EWSR1, we applied the H *(r)* transformation of Ripley’s *k*-statistic to whole nuclei STED images. The Ripley’s *k*-statistic and its derivatives evaluate relative clustering as a function of spatial distance.^30–32^ Representative analyses from HT-1080-FF-EWSR1 cells and the other FF-EWSR1 cell lines showed a leftward shift of the frequency curves relative to a random organization (dashed line) (**Figures 1H** and **S2H**). These results are consistent with endogenous EWSR1 exhibiting a non-random spatial clustering reflective of a highly organized nuclear architecture.

### EWSR1 and newly synthesized RNA form a ribonucleoprotein network

To further examine endogenous EWSR1’s nuclear organization, we examined its spatial distribution relative to nucleic acids. EWSR1 contains two nucleic acid-binding domains, a zinc finger and an RRM. Using A673- and TC-32-mNG-EWSR1 reporters stained with DAPI, we previously showed at ∼120 nm resolution that mNG-EWSR1 and DNA exhibit distinct non-overlapping signals and the exclusion of mNG-EWSR1 from regions of condensed DNA. In contrast, mNG-EWSR1 exhibited close spatial organization with newly synthesized RNA throughout the nucleoplasm. To extend these observations, we first analyzed the FF-EWSR1 reporter cell lines employing anti-FLAG IF in combination with the SiR-DNA dye, which together with STED microscopy enabled direct comparison of EWSR1 and DNA localization at an increased spatial resolution.

Consistent with our previous observations^26^, we observed DNA and FF-EWSR1 in proximity to each other within the nucleus (**Figures 2A** and **2B**, **Figures S3A–D**). To quantitatively assess the relative spatial organization of FF-EWSRI and DNA, we applied an adaptation of Ripley’s *k* function to first define the point pattern distribution of SiR-DNA signals, generating DNA reference structures for each FF-EWSR1 reporter cell line. Subsequently, we evaluated the spatial distribution of FF-EWSR1 relative to these cell line specific DNA reference patterns. Cumulative frequency plots for FF-EWSR1 and DNA from four nuclei per FF-EWSR1 reporter cell line showed that the majority of FF-EWSR1 exhibits spatial clustering with DNA, while a distinct fraction of FF-EWSR1 displayed a distribution indistinguishable from random with respect to DNA (**Figures 2C** and **S3E).** These data suggested that DNA could contribute to the spatial organization of a subset of EWSR1.

**Figure 2:**
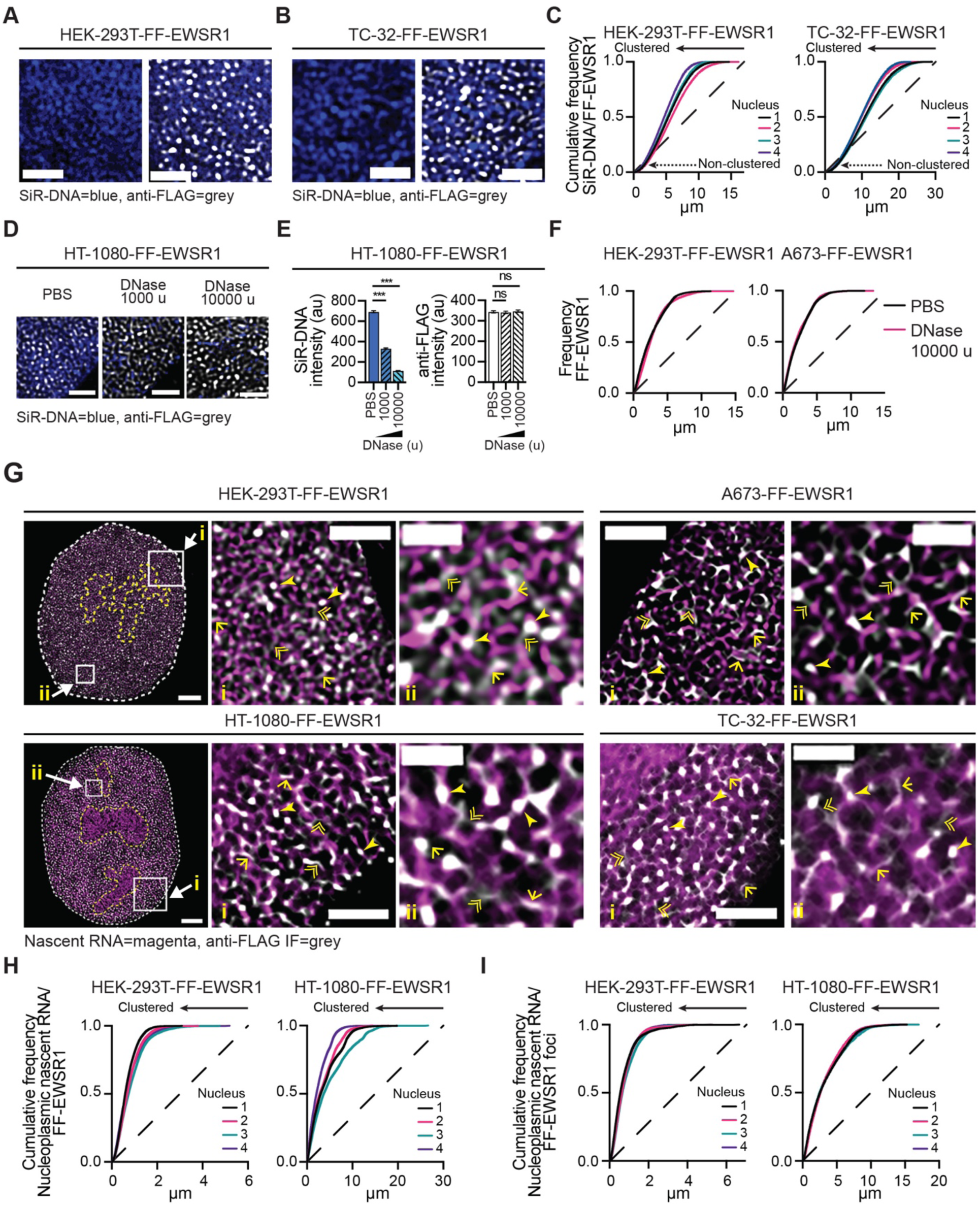
EWSR1 and newly synthesized RNA form a ribonucleoprotein network. (**A** and **B**) STED microscopy images of expanded nuclear regions from HEK-293T-FF-EWSR1 (**A**) and TC-32-FF-EWSR1 (**B**) cells. Single-channel images show SiR-DNA (blue), and merged images show SiR-DNA (blue) and anti-FLAG IF (grey). Corresponding nuclei images with expanded regions indicated are shown in **Figures S3A** and **S3B**. Scale bar, 1 µm. (**C**) Spatial cluster analysis of FF-EWSR1 IF pattern distribution relative to DNA, using SiR-DNA as the reference structure. Cumulative frequency plots show analysis of STED microscopy images of four representative nuclei per indicated cell line. (**D**) STED microscopy images of expanded nuclear regions from HT-1080-FF-EWSR1 cells treated with either PBS, or DNase (1000 or 10000 units (u)) for 20 min. Merged images show anti-FLAG IF (grey) and SiR-DNA (blue). Corresponding nuclei images are shown in **Figure S3F**. Scale bar, 1 µm. (**E**) Quantification of the SiR-DNA and anti-FLAG fluorescence intensity in HT-1080-FF-EWSR1 reporter cell lines following treatment with PBS or DNase (1000 or 10000 u, 20 min). (**F**) Spatial cluster analysis of FF-EWSR1 IF pattern distribution. Plots show analysis of STED microscopy images of representative nuclei from HEK-293T-FF-EWSR1 and A673-FF-EWSR1 cells treated with PBS or DNase (10000 u). (**G**) STED microscopy images of nuclei and expanded nuclear regions (i and ii) from HEK-293T-FF-EWSR1 and HT-1080-FF-EWSR1 cells and expanded nuclear regions (i and ii) from A673-FF-EWSR1 and TC-32-FF-EWSR1 cells (corresponding nuclei images are shown in **Figure S4A**). Merged images show anti-FLAG IF (grey) and nascent RNA labeling (magenta). Single arrowheads indicate high-intensity FF-EWSR1 foci colocalized with RNA, single arrowed lines indicate lower-intensity distributed FF–EWSR1 signals close to RNA, double arrowheads indicate lower-intensity distributed FF–EWSR1 signal. Scale bars, nucleus, 2 µm, expanded regions i, 1 µm, ii, 0.5 µm. (**H, I**) Spatial cluster analysis of FF-EWSR1 IF (**H**) and high intensity (≥5x signal) FF-EWSR1 foci (**I**) pattern distribution relative to newly synthesized RNA in the indicated FF-EWSR1 reporter cell lines using EU-labeled RNA as the reference structure. Cumulative frequency plots show analysis of STED microscopy images of four representative nuclei per indicated cell line. (**A**, **B**, **D**, **G**) Images shown are representative of >20 nuclei per indicated FF-EWSR1 reporter cell line. (**C**, **F, H, I**) The dashed line indicates a random point pattern distribution; arrowed solid lines indicate deviations from a random point pattern distribution (**C**, **H, I**); arrowed dotted lines indicate non-clustered signals (**C**). (**E**) Data are presented as mean ± SEM of 20 nuclei per treatment per reporter cell line. Statistical significance was determined by one-way ANOVA, _***_ p<0.001, ns non-significant.

To test this hypothesis, we examined the effect of DNA degradation on EWSR1. Representative STED images of nuclei and expanded regions from FF-EWSR1 cell lines treated with either vehicle (PBS), or DNase (1000 or 10000 units (u) for 20 minutes (min)) are shown in **Figure 2D** and **Figures S3F** and **S3G**. Quantification of the anti-FLAG IF and SiR-DNA signals showed that despite a concentration-dependent reduction in SiR-DNA fluorescence following DNase treatment, there was no corresponding change in the anti-FLAG IF signal (**Figures 2E** and **S3H**). To determine whether DNase-mediated DNA degradation altered EWSR1 organization, we quantified the clustering of anti-FLAG IF signals across nuclei from each FF-EWSR1 reporter cell line treated with PBS or 10000 u DNase (20 min) (**Figures 2F** and **S3I**). DNase-mediated DNA degradation did not produce any change in FF-EWSR1 spatial organization, indicating that the nuclear distribution of EWSR1 is independent of DNA.

We next examined the spatial relationship between EWSR1 and RNA. To selectively visualize newly synthesized RNA, we used 5-ethynyl uridine (EU) incorporation for 40 min followed by Click-iT conjugation to Alexa Fluor 594. Representative merged STED images of FF-EWSR1 reporter cell lines show extensive spatial proximity of EWSR1 (anti-FLAG IF) and EU-labelled nascent RNA (magenta) (**Figures 2G** and **S4A**). STED microscopy, at a resolution of <50 nm, shows endogenous EWSR1 and newly synthesized RNA exhibit a network-like organization. Specifically, EWSR1 foci extensively overlap with nascent RNA signals (single arrowheads) to form the nodes of a network, with distributed EWSR1 (lower intensity signals) linking many of these nodes (double arrowheads), which in some cases align with EU-labelled nascent RNA (single arrowed lines).

To assess this network-like organization using statistical analysis, we quantified the clustering of EWSR1 and newly synthesized RNA by applying the adaptation of the Ripley’s *k* function to define the cell line-specific point pattern distribution of nucleoplasmic nascent RNA and assessed the spatial organization of FF-EWSR1 relative to these reference structures. Cumulative frequency plots from four nuclei per FF-EWSR1 reporter cell line showed significant clustering of FF-EWSR1 in the context of nucleoplasmic nascent RNA (**Figures 2H** and **S4B**). This clustering of EWSR1 with RNA contrasts with its partially random point pattern distribution seen in the context of DNA (**Figures 2C** and **S3E**). Next, we examined the clustering of EWSR1 foci relative to nucleoplasmic nascent RNA. We focused on analysis of EWSR1 foci as the relative contribution of distributed EWSR1 alone cannot be statistically isolated, but threshold-based segmentation enabled selective analysis of the high intensity EWSR1 focal signals. In prior analysis using A673- and TC-32-mNG-EWSR1 cells, we defined foci as regions exhibiting anti-FLAG fluorescence intensity ≥3x over background^26^. Given the increased spatial resolution afforded by STED microscopy, we applied a more stringent criterion here, defining EWSR1 foci as regions with anti-FLAG fluorescence intensity ≥5x over background (**Figure S4C**).

Applying the ≥5x background fluorescence threshold to central plane images from 30 nuclei per FF-EWSR1 cell lines, we quantified the number and diameter of EWSR1 foci (**Figure S4D**). Three of the four cell lines exhibited comparable numbers of FF-EWSR1 foci per central z-plane. TC-32 cells had slightly more foci, but this cell line also had larger nuclei compared with the others (nuclei diameter, mean ± SEM (n=100, per cell line) HEK-293T 7.84±0.22 µm, HT-1080 18.9±0.34 µm, A673 9.89±0.28 µm, TC-32 22.5±0.40 µm, **Figure S4E**). In addition, both A673 and TC-32 EWS cell lines exhibited slightly larger FF-EWSR1 foci, which, we propose reflects the enhanced transcriptional activity promoted by the presence of the EWSR1::FLI1 fusion oncoprotein in these cell lines.^33–36^ Nevertheless, irrespective of the cell line, cumulative frequency analyses demonstrated significant clustering of FF-EWSR1 foci relative to nucleoplasmic nascent RNA (**Figures 2I** and **S4F**), indicating an association between EWSR1 foci and active RNA synthesis.

Taken together, the images of EWSR1 and newly synthesized RNA, along with quantitative analyses of their point pattern distributions (**Figures 1, 2**, **S2** and **S4**), show that EWSR1 and nascent RNA exhibit an interconnected organization consistent with the formation of an EWSR1-centered RNP network.

### The depletion of EWSR1 results in a transient decrease in newly synthesized RNA and cell viability

To determine whether formation of the EWSR1-RNP network represents a fundamental function of EWSR1, we acutely depleted endogenous EWSR1 using the FKBP12^F36V^/dTAG-13 degron system (**Figure S5A**). Representative merged STED images of TC-32-FF-EWSR1 cells treated with vehicle (DMSO) or dTAG-13 (50 or 5000 nM) for 0.5 or 2 hours (hrs) are shown in **Figure 3A**, with corresponding images from similarly treated HT-1080-FF-EWSR1 cells shown in **Figure S5B**.

**Figure 3:**
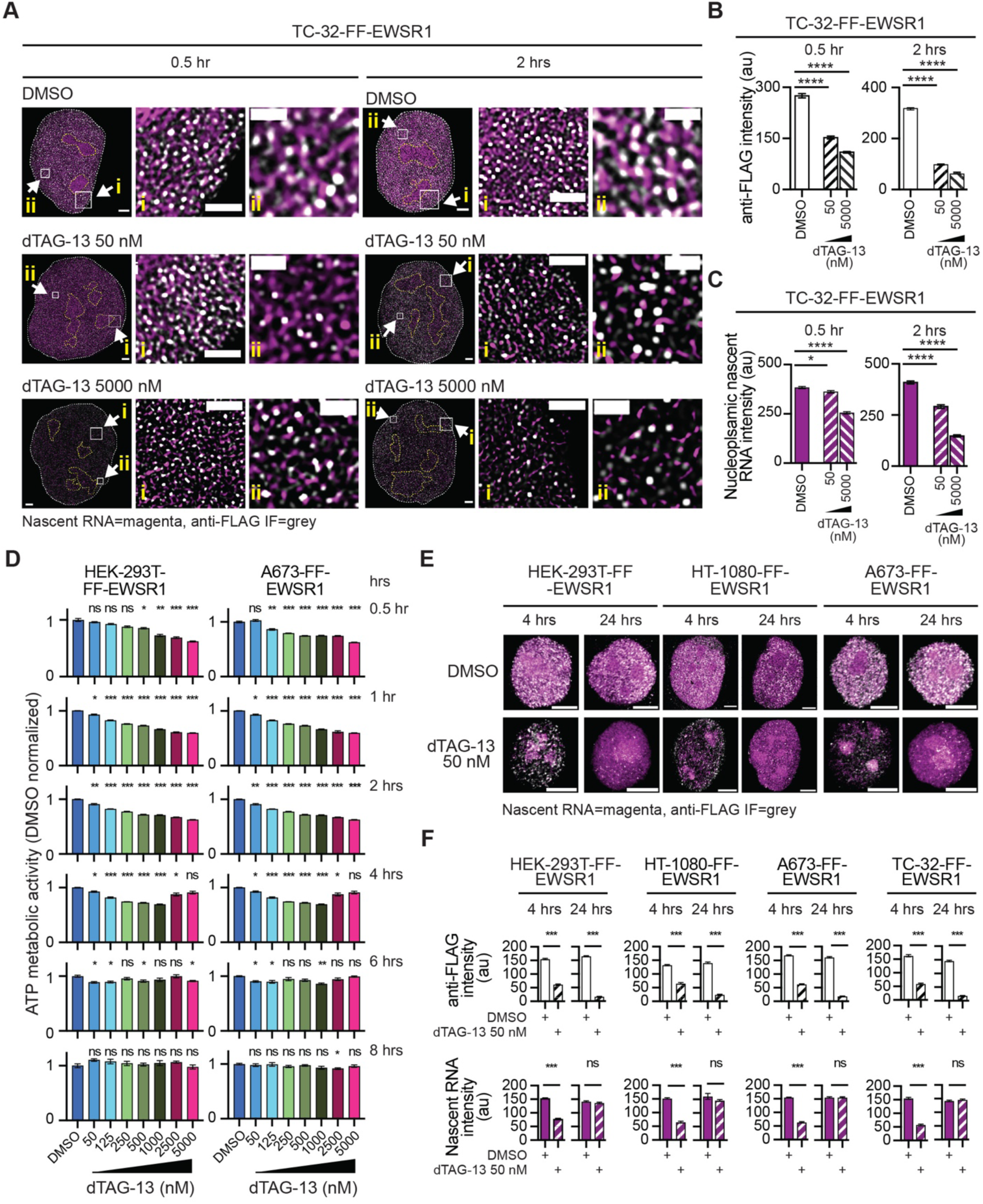
The degradation of EWSR1 results in a transient decrease in nascent RNA and cell viability. (**A**) STED microscopy images of nuclei and two expanded nuclear regions (i and ii) from TC-32-FF-EWSR1 cells treated with DMSO (upper panels), or 50 nM (middle panels) or 5000 nM (lower panels) dTAG-13 for 0.5 hrs (left) or 2 hrs (right). Merged images show anti-FLAG IF (grey) and nascent RNA (magenta). Scale bars, nucleus, 2 µm, expanded regions i, 1 µm, ii, 0.5 µm. (**B, C**) Quantification of anti-FLAG IF intensity (**B**) and nascent RNA fluorescence intensity (**C**) in TC-32-FF-EWSR1 cells following treatment with DMSO or dTAG-13 (50 nM or 5000 nM) for 0.5 hrs (left) or 2 hrs (right). (**D**) Quantification of ATP metabolic activity in HEK-293T-FF-EWSR1 and A673-FF-EWSR1 reporter cell lines measured 0.5 – 8 hrs after addition of DMSO or increasing concentrations of dTAG-13 (50 - 5000 nM). Data were normalized to the mean viability of DMSO-treated cells at each time point. (**E**) SoRa images of nuclei from the indicated FF-EWSR1 reporter cell lines 4 or 24 hrs after treatment with DMSO or dTAG-13 (50 nM). Merged images show anti-FLAG IF (grey) and nascent RNA (magenta). Scale bars, 4 µm. (**F**) Quantification of anti-FLAG IF intensity (upper graphs) and nascent RNA fluorescence intensity (lower graphs) in the indicated FF-EWSR1 reporter cell lines 4 or 24 hrs after treatment with DMSO or dTAG-13 (50 nM). (**A**, **E**) Images shown are representative of >20 nuclei from the indicated FF-EWSR1 cell line. (**B**-**D**, **F**) Data shown as mean ± SEM. (**B** and **C**) n=15 nuclei per cell line, (**D**) n= 6 biological replicates per cell line per treatment per time point. (**F**) n=20 nuclei per cell line per condition. (**B** - **D**) Statistical significance was determined using one-way ANOVA. (**F**) Statistical significance was determined using Welch’s t test. (**B, C, D** and **F**) _*_ p<0.05, _***_ p<0.001, _****_ p<0.0001, ns non-significant.

Quantification of anti-FLAG IF intensities revealed a concentration- and time-dependent reduction in EWSR1 signal, consistent with the selective degradation of endogenous EWSR1 (**Figures 3B** and **S5C**). In parallel, following 2 hrs exposure to dTAG-13, there was a modest change in the number and diameter of FF-EWSR1 foci, particularly in TC-32-FF-EWSR1 cells (**Figure S5D**). Strikingly, global reduction in nascent RNA signal, as measured by EU incorporation accompanied the acute depletion of EWSR1 in both EWS (TC-32) and non-EWS (HT-1080) cell lines (**Figures 3C** and **S5E**). These findings indicate that loss of EWSR1 rapidly compromises RNA synthesis and/or accumulation, supporting a direct role for EWSR1 in sustaining global gene expression.

A reduction in the synthesis or accumulation of pre-mRNA could impair multiple energy-dependent processes. We thus next asked whether acute degradation of EWSR1 affects cellular metabolic activity of cells. Using an ATP-based assay, we assessed metabolic activity in the FF-EWSR1 reporter cell lines treated with increasing concentrations of dTAG-13 (50 – 5000 nM) for 0.5 – 8 hrs (**Figures 3D** and **S5F**). At 0.5 hrs, the highest dTAG-13 concentrations (2,500 and 5000 nM) caused a significant reduction in ATP levels compared to DMSO-treated cells. By 1 – 2 hrs, metabolic activity decreased in a concentration-dependent manner at all dTAG-13 concentrations. Notably, after 4 hrs of treatment, ATP levels remained suppressed at lower dTAG-13 concentrations but partially rebounded at the two highest concentrations. By 6 hrs, metabolic activity had returned to near baseline levels at most concentrations, and by 8 hrs, we detected no significant differences in ATP levels at any dTAG-13 concentration. Together, these data indicate that EWSR1 depletion induces a rapid but transient reduction in cellular metabolic activity consistent with the presence and activation of compensatory mechanisms that restores cellular function despite sustained loss of EWSR1.

To assess whether the recovery of metabolic activity observed following EWSR1 depletion is associated with restoration of RNA synthesis or accumulation, we imaged FF-EWSR1 reporter cell lines treated with 50 nM dTAG-13, the lowest dTAG-13 concentration used in the assay of metabolic activity, for 4 or 24 hrs (**Figures 3E** and **S5G**) and quantified anti-FLAG IF and nascent RNA fluorescence. Despite sustained depletion of EWSR1, the decrease in nascent RNA was transient, with complete recovery observed by 24 hrs (**Figure 3F**). These data indicate that degrading EWSR1 results in an acute reduction in nascent RNA and that this is associated with a decrease in metabolic activity. However, there is a compensatory mechanism that restores the synthesis and/or accumulation of newly synthesized RNA and reverses the deleterious effects on cell metabolism.

Together, these results support a model in which EWSR1 supports the maintenance of nascent RNA levels under basal conditions, while adaptive responses can substitute for its loss.

### EWSR1 foci colocalize with actively elongating RNA polymerase II, but the depletion of EWSR1 does not measurably alter this marker of active transcription

To elucidate the basis for the rapid reduction in nascent RNA and cellular metabolic activity following EWSR1 degradation, we asked whether the biochemical interaction between EWSR1 and RNA pol II reflects an EWSR1-dependent role in transcription. Consistent with EWSR1 functioning as a component of the transcriptional machinery, we previously showed that EWSR1 foci significantly colocalize with pS5- and pS2-RNA pol II in A673-mNG-EWSR1 and TC-32-mNG-EWSR1 cells^26^. To confirm and extend these observations at higher spatial resolution, we examined the localization of FF-EWSR1 relative to transcriptionally active RNA pol II using STED microscopy and detection of pS2-RNA poll II as a marker of elongating polymerase (**Figure 4A**).

**Figure 4:**
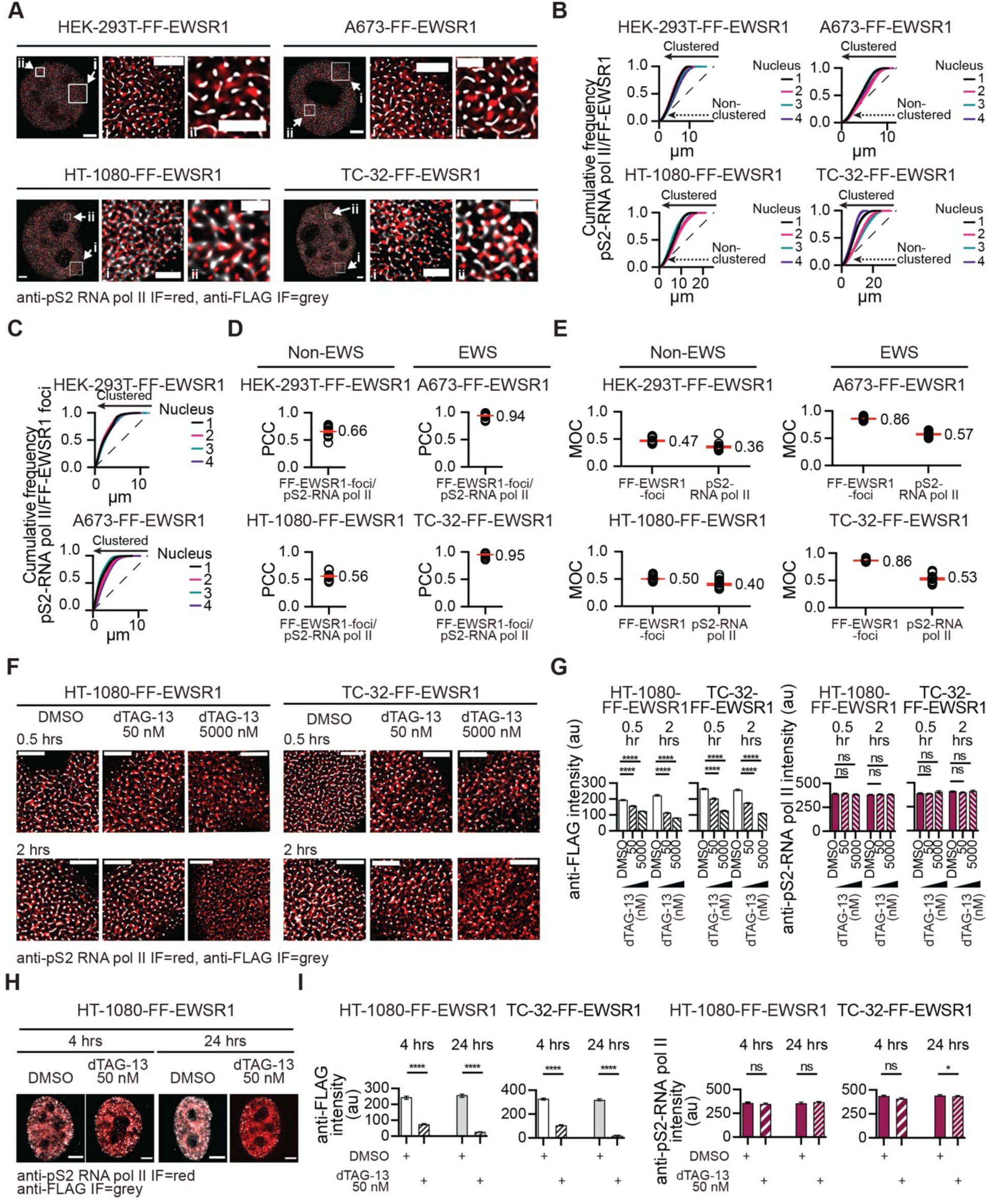
EWSR1 foci colocalize with actively elongating RNA polymerase II, but depletion of EWSR1 does not measurably alter this marker of transcriptional activity. (**A**) STED microscopy images of nuclei and expanded nuclear regions (i and ii) from the indicated FF-EWSR1 reporter cell lines. Merged images show anti-FLAG IF (grey) and pS2 RNA pol II IF (red). Scale bars, nucleus, 2 µm, expanded regions i, 1 µm, ii, 0.5 µm. (**B, C**) Spatial cluster analysis of FF-EWSR1 (**B**) and FF–EWSR1 foci (≥5x above background) (**C**) pattern distributions using pS2-RNA pol II IF as the reference structure in the indicated FF–EWSR1 reporter cell lines. Cumulative frequency plots generated using STED microscopy images of four representative nuclei per indicated cell line per analysis. (**D, E**) Quantification of colocalization between FF-EWSR1-foci and pS2 RNA pol II in the indicated FF-EWSR1 reporter cell lines using Pearson correlation coefficient (PCC) (**D**) and Mander’s overlap coefficient (MOC) (**E**) analysis. (**F**) STED microscopy images of expanded nuclear regions from HT-1080-FF-EWSR1 and TC-32-FF-EWSR1 cells treated with DMSO or dTAG-13 (50 or 5000 nM) for the indicated times. Merged mages show anti-FLAG (grey) and pS2-RNA pol II IF (red). Corresponding whole-nucleus images are shown in **Figure S6C, D**. Scale bar, 1 µm. (**G**) Quantification of anti-FLAG IF (left) and pS2-RNA pol II IF (right) intensities in the indicated FF-EWSR1 reporter cell lines following treatment with DMSO or dTAG-13 (50 nM or 5000 nM) for 0.5 or 2 hrs. (**H**) SoRa super resolution images of nuclei from the indicated FF-EWSR1 reporter cell lines 4 or 24 hrs after treatment with DMSO or dTAG-13 (50 nM). Merged mages show anti-FLAG (grey) and pS2-RNA pol II IF (red). Scale bar, 4 µm. (**I**) Quantification of anti-FLAG IF intensity (left) and pS2-RNA pol II IF intensity (right) in the indicated FF-EWSR1 reporter cell lines treated with DMSO or dTAG-13 (50 nM) for 4 or 24 hrs. (**A**, **F, H**) Images are representative of >20 nuclei per indicated FF-EWSR1 reporter cell line. (**B, C**) The dashed line represents a random point pattern distribution; arrowed solid lines indicate deviations from a random point pattern distribution; arrowed dotted lines indicate non-clustered signals (**B**). (**D-E**) Data are shown as individual data points and mean ± SEM (red lines, mean value indicated), n=20 nuclei. (**G, I**) Data are shown as mean ± SEM, n=20 nuclei per treatment per time point. Statistical significance was determined using one-way ANOVA (**G**) and Welch’s t test (**I**). (**G** and **I**) _*_ p<0.05, _***_ p<0.00, _****_ p<0.0001, ns non-significant.

Using whole nucleus images (four nuclei per cell line), we applied Ripley’s *k* function analysis to define the point pattern distribution of pS2-RNA pol II to generate cell line-specific reference structures. We next quantified the spatial clustering of FF-EWSR1 relative to these reference patterns. Cumulative frequency analyses revealed that a substantial proportion of FF-EWSR1 signals cluster with pS2-RNA pol II signals, but not all (**Figure 4B**). To assess whether the EWSR1 signals that cluster with pS2-RNA pol II reflect the colocalization of EWSR1 foci with phosphorylated RNA pol II we observed previously^26^, we stringently defined FF-EWSR1 foci using a ≥5x background fluorescence intensity threshold (**Figure S6A**). Under these conditions, cumulative frequency plots demonstrated near-complete clustering of FF-EWSR1 foci with pS2-RNA pol II fluorescence (**Figures 4C** and **S6B**). These results indicate that FF-EWSR1 foci represent localized concentration of much of the EWSR1 protein at sites of active transcription.

To further quantify the spatial proximity between FF-EWSR1 foci and actively elongating RNA pol II, we analyzed images from 20 nuclei per cell line and calculated both the Pearson correlation coefficient (PCC) (**Figure 4D**) and the Mander’s overlap coefficient (MOC) (**Figure 4E**) as complementary measures of their colocalization.^37,38^ PCC analysis revealed substantial colocalization of FF-EWSR1 foci and pS2-RNA pol II in non-EWS cell lines (PCC ∼0.6), with markedly higher colocalization in EWS cell lines (PCC >0.9). MOC analysis also demonstrated significantly greater overlap between FF-EWSR1 foci and pS2-RNA pol II in EWS cells (FF-EWSR1 foci MOC ∼0.86) compared with non-EWS cells (FF-EWSR1 foci MOC ∼0.5). Reciprocal MOC measurements further indicated that a larger fraction of elongating RNA pol II overlapped with FF-EWSR1 foci in EWS cells (pS2-RNA pol II MOC ∼0.55) than in non-EWS cells (pS2-RNA Pol II MOC ∼0.4). Together, these quantitative analyses support an enhanced association between EWSR1 and transcriptionally active polymerase in the EWS cells when compared to non-EWS cells.

The increased colocalization of EWSR1 with transcriptionally active pS2-RNA pol II observed in EWS cells is consistent with the activity of the EWSR1::FLI1 fusion oncoprotein, which extensively reprograms gene expression.^39^ One consequence of this transcriptional reprogramming could be the enhanced association of EWSR1 with sites of active transcription. In addition, or alternatively, the EWSR1::FLI1 fusion oncoprotein may modify the interaction of EWSR1 with the RNA pol II machinery, either directly or through changes in transcriptional complex composition. Importantly, despite quantitative differences between non-EWS and EWS cell lines, our analyses demonstrate that within the nucleoplasm, most EWSR1 foci and a substantial fraction of transcriptionally active RNA pol II colocalize. Based on these observations, we next examined whether depletion of EWSR1 alters transcriptional activity as measured by anti-pS2-RNA pol II IF.

To determine whether loss of EWSR1 affects transcriptionally active RNA pol II, we examined merged anti-pS2-RNA pol II and anti-FLAG IF images of HT-1080- and TC-32-FF-EWSR1 cells treated with either vehicle (DMSO) or dTAG-13 (50 or 5000 nM) for 0.5 or 2 hrs (**Figures 4F** and **S6C, D**) and quantified their respective fluorescence intensities under each condition (**Figure 4G**). Despite the pronounced reductions in nascent RNA and metabolic activity observed at 0.5 or 2 hrs post-addition of dTAG-13 (**Figures 3A-C** and **S5B-D**), pS2 RNA pol II fluorescence intensities were unchanged. Immunoblot analysis of whole cell lysates prepared from TC-32-FF-EWSR1 cells treated with DMSO or dTAG-13 for 0.5 and 2 hrs also showed no changes in pS2-RNA pol II (**Figure S6E**). We next extended this analysis to later time points and found that treatment of HT-1080- and TC-32-FF-EWSR1 cells with 50 nM dTAG-13 for 4 or 24 hrs did not alter anti–pS2-RNA pol II fluorescence intensity at 4 hrs, and resulted in only a modest decrease at 24 hrs in TC-32-FF-EWSR1 cells (**Figures 4H**, **I** and **S6F**).

Although pS2–RNA pol II captures only one aspect of the transcriptional process, these data suggest that global changes in transcriptionally active RNA pol II do not account for either the acute loss or the subsequent recovery of newly synthesized RNA observed following EWSR1 degradation. Nevertheless, the concentration of EWSR1 at RNA pol II is consistent with a role for EWSR1 in maintaining RNA levels at sites of active transcription and that the recovery of newly synthesized RNA following EWSR1 degradation for greater than 4 hrs involves a compensatory mechanism in which another protein functionally substitutes for EWSR1.

### FUS and TAF15 functionally compensate for EWSR1 loss to restore cellular metabolic activity

The EWSR1 homologs FUS and TAF15 share highly similar domain architectures with EWSR1, including conserved nucleic acid–binding motifs and multiple LCDs (**Figure 5A**) raising the possibility of functional redundancy. To test whether FUS or TAF15 can compensate for EWSR1 loss, we first examined the combined effects of their depletion on cellular metabolic activity using a luminescence-based ATP assay. FF-EWSR1 cells were untreated or transfected with a control siRNA (siNeg), an siRNA targeting *FUS* (siFUS; **Figure S7A**), an siRNA targeting *TAF15* (siTAF15; **Figure S7B**), or both siFUS and siTAF15. Forty-eight hours post-transfection, we treated cells with DMSO or dTAG-13 (5000 nM) for 2, 8, or 24 hrs (**Figures 5B, C** and **S7C**).

**Figure 5:**
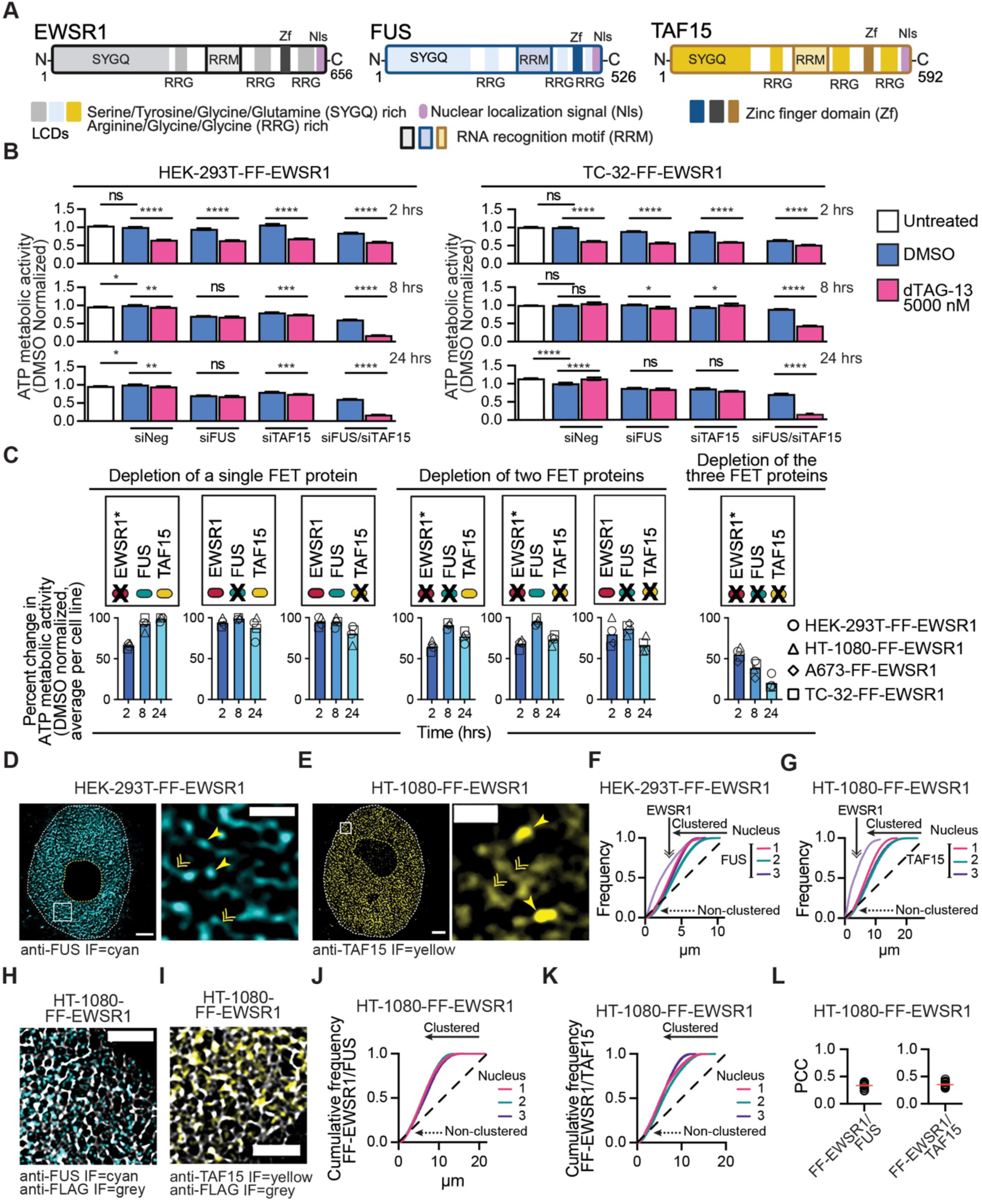
FUS and TAF15 functionally compensate for EWSR1 loss to restore cellular metabolic activity. (A) Organization of EWSR1, FUS, and TAF15 highlighting their common organizational features, specifically, nucleic acid binding (RRM, Zf), low-complexity (SYGQ, RGG), and C-terminal nuclear localization signal (Nls) domains. (B) The ATP metabolic activity of HEK-293T-FF-EWSR1 and TC-32-FF-EWSR1 reporter cell lines following siRNA-mediated depletion of FUS and/or TAF15 (48 hrs) and dTAG-13-mediated degradation of EWSR1 (2, 8, or 24 hrs). Data are normalized to the mean of the siNeg-transfected, DMSO-treated cells. (C) Summary of ATP metabolic activity across FF-EWSR1 reporter cell lines highlighting rescue by FUS and TAF15. Schematics indicate the depleted protein (indicated by X) and the mode of depletion (*dTAG-13 for EWSR1; RNAi for *FUS* and *TAF15*). (D) STED microscopy images of a nucleus and an expanded nuclear region from a HEK-293T-FF-EWSR1 showing FUS IF (cyan). The double arrowheads indicate lower intensity fluorescence or distributed FUS, and the single arrowheads indicate higher intensity fluorescence or FUS foci. Scale bars, nucleus, 2 µm, expanded region, 1 µm. (E) STED microscopy images of a nucleus and an expanded nuclear region from a HT-1080-FF-EWSR1 showing TAF15 IF (yellow). The double arrowheads indicate lower intensity fluorescence or distributed TAF15, and the single arrowheads indicate higher intensity fluorescence or TAF15 foci. Scale bars, nucleus, 2 µm, expanded region, 1 µm. (F) Spatial cluster analysis of FUS IF pattern distribution for the HEK-293T-FF-EWSR1 reporter cell line. Frequency plots generated using STED microscopy images of three nuclei. A frequency plot of FF-EWSR1 in the same cell line is shown for comparison (light purple). (G) Spatial cluster analysis of TAF15 IF pattern distribution for the HT-1080-FF-EWSR1 reporter cell line. Frequency plots generated using STED microscopy images of three nuclei. A frequency plot of FF-EWSR1 in the same cell line is shown for comparison (light purple). (**H, I**) STED microscopy images of expanded regions of HT-1080-FF-EWSR1 nuclei showing anti-FUS (cyan) and anti-FLAG IF (grey)(**H**) and anti-TAF15 IF (yellow) and anti-FLAG IF (grey)(**I**) . Scale bar, 1 µm. Corresponding nuclei images are shown in **Figure S8C, 8D**. (**J, K**) Spatial cluster analysis of FUS IF (**J**) and TAF15 IF (**K**) pattern distributions for the HT-1080-FF-EWSR1 reporter cell line using FF-EWSR1as the reference structures. Cumulative frequency plots show analysis of the STED images of three nuclei per analysis. (**L**) Quantification of FF–EWSR1 colocalization with FUS or TAF15 by PCC analysis. (**D, E, H, I**) Images are representative of >20 nuclei per the indicated FF-EWSR1 reporter cell line. (**B**) Data are shown as mean ± SEM, n=6 per treatment per time point. Statistical significance was determined using one-way ANOVA. _*_ p<0.05, _**_ p<0.01, _***_ p<0.001, _****_ p<0.0001, ns non-significant. (**F, G, J, K**) The dashed line represents a random point pattern distribution; arrowed solid lines indicate deviations from a random point pattern distribution; arrowed dotted lines indicate non-clustered signals. (**L**) Data are shown as individual values and mean ± SEM (red lines), n=20 nuclei per assessment. Schematics in **(A)** and **(C)** were generated using BioRender.

At 2 hrs following dTAG-13 treatment, all cells exhibited a comparable reduction in metabolic activity relative to DMSO-treated control cells, regardless of FUS or TAF15 expression status, indicating that the acute metabolic effects of EWSR1 degradation are independent of these homologs. In contrast, at 8 and 24 hrs post-dTAG-13 treatment, cells retaining FUS or TAF15 consistently displayed higher metabolic activity than at 2 hrs post addition of dTAG-13. However, the combined depletion of all three FET family proteins, EWSR1 via targeted degradation and FUS and TAF15 via RNAi, resulted in a pronounced and sustained reduction in metabolic activity at these time points. To strengthen these findings, we assessed metabolic activity in HT-1080-FF-EWSR1 and HEK-293T-FF-EWSR1 cells stably expressing an *EWSR1* cDNA. In addition to siRNAs targeting *FUS*, *TAF15*, or both, we included a condition combining siFUS and siTAF15 with an siRNA targeting the *EWSR1* coding region (siEWSR1 s4888; **Figure S1E**). Following siRNA transfection, we treated cells with dTAG-13 (5000 nM) to induce EWSR1 degradation and measured metabolic activity 2, 8, or 24 hrs later. Expression of the *EWSR1* cDNA rescued the effects of EWSR1 degradation on metabolic activity under all conditions except when *FUS*, *TAF15*, and *EWSR1* were simultaneously silenced (**Figure S7D**). Together, these results indicate that cellular metabolic homeostasis requires the presence of at least one FET family protein and that FUS and TAF15 can compensate for EWSR1-dependent functions underlying the acute metabolic defects observed upon EWSR1 degradation.

To further investigate the compensatory mechanism revealed by EWSR1 degradation, we next asked whether FUS and TAF15 exhibit nuclear organizations comparable to that of EWSR1. STED microscopy of whole nuclei and corresponding expanded regions from HEK-293T-FF-EWSR1 and TC-32-FF-EWSR1 cells show widespread distribution of FUS (anti-FUS IF, cyan) throughout the nucleoplasm (**Figures 5D** and **S7E**). Similarly, STED imaging of HT-1080-FF-EWSR1 and A673-FF-EWSR1 cells showed that TAF15 (anti-TAF15, yellow) displays a comparable nucleoplasmic localization (**Figures 5E** and **S7F**). Like EWSR1, FUS and TAF15 exhibited two distinct fluorescent signals, characterized by regions of lower intensity, diffuse IF signals (distributed) and discrete higher intensity puncta consistent with the formation of foci. To quantitatively characterize this organization, we applied a ≥5x background fluorescence intensity threshold to define FUS and TAF15 foci and assessed their number and diameter in the central z-plane of 20 nuclei per cell line (**Figures S7G, H**). Across cell lines, EWS cells had modestly fewer FUS and TAF15 foci than non-EWS cells. Interestingly, FUS foci were smaller in diameter than TAF15 and EWSR1 foci (foci diameter: FUS ∼70 nm, TAF15 ∼150 nm, EWSR1 >100 nm, **Figure S6A**).

To directly compare the spatial organization of FUS and TAF15 with that of EWSR1, we analyzed anti-FUS and anti-TAF15 whole nucleus STED images (three nuclei per cell line) and quantified the clustering of above-background fluorescence intensities (**Figures 5F, G** and **S8A, B**). These analyses revealed that although both FUS and TAF15 display substantial clustering within the nucleoplasm, the degree of clustering is less than that obtained for EWSR1.

We next examined the organization of FUS and TAF15 relative to EWSR1. **Figures 5H**, **I** and **Figures S8C-F**, show representative STED images of nuclei and expanded regions from HT-1080-FF-EWSR1 and TC-32-FF-EWSR1 cells probed for FUS and FF-EWSR1 or TAF15 and FF-EWSR1. To assess the organization of FUS and TAF15 relative to EWSR1, we determined the point pattern distribution of FF-EWSR1 in HT-1080-FF-EWSR1 and TC-32-FF-EWSR1 cells to generate cell-line specific reference structures and compared the distribution of FUS and TAF15 to these reference structures. The cumulative frequency curves suggest that while a fraction of FF-EWSR1 clusters with either FUS or TAF15, a proportion of FF-EWSR1 does not cluster with either homolog (**Figures 5J, K** and **S8G, H**). We confirmed this conclusion by quantifying colocalization using PCC and MOC metrics for FF-EWSR1–FUS and FF-EWSR1–TAF15 pairs (**Figures 5L** and **S8I-K**). Analysis of images from both cell lines calculated mean PCC values consistently below 0.4 and mean MOC values of less than 0.5 for FF-EWSR1/FUS and FF-EWSR1/TAF15. The minimal overlap of FUS and TAF15 with FF-EWSR1 under unperturbed conditions led us to hypothesize that, upon EWSR1 loss, changes in the expression level and/or nuclear organization of FUS and TAF15 enable them to compensate for EWSR1-dependent functions.

### FUS and TAF15 reorganize following EWSR1 degradation

All three FET family proteins contain a conserved 87–amino acid RRM. The RRMs of FUS and TAF15 are similar (72% identity), while EWSR1’s RRM shares 60% and 59% identity with those of FUS and TAF15, respectively (**Figure S9A**). Given our observation that EWSR1 exhibits significant spatial clustering with nucleoplasmic nascent RNA (**Figures 2I** and **S4F**), we next asked whether FUS and TAF15 display comparable associations with newly synthesized RNA (**Figures 6A** and **6B**). To quantify the spatial relationship between FUS or TAF15 and nascent RNA, we generated cell line–specific point-pattern reference structures for nucleoplasmic nascent RNA and analyzed the distribution of FUS and TAF15 relative to these patterns. Both FUS and TAF15 exhibited clustering with nucleoplasmic nascent RNA (**Figures 6C** and **6D**). However, in contrast to EWSR1 (**Figures 2I** and **S4F**), a fraction of the FUS and TAF15 signals did not cluster with EU-labeled RNA (**Figures 6C** and **6D**, dashed arrowed lines), indicating a more heterogeneous association with newly synthesized RNA than that of EWSR1.

**Figure 6:**
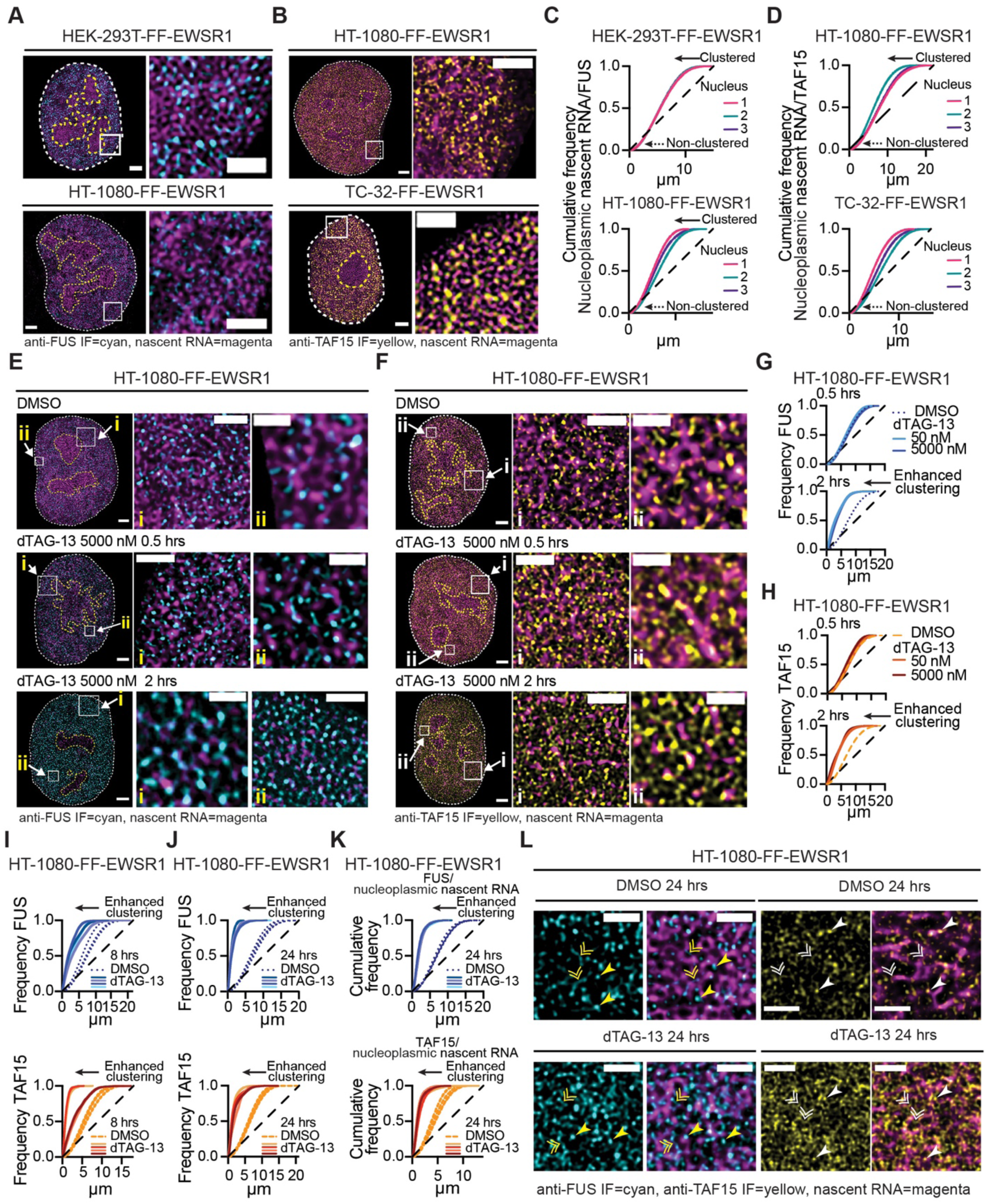
FUS and TAF15 reorganize following EWSR1 degradation. (**A**) STED microscopy images of nuclei and expanded nuclear regions from HEK-293T-FF-EWSR1 (upper) and HT-1080-FF-EWSR1 (lower) cells. Merged images show anti-FUS (cyan) and nascent RNA (magenta). Scale bars, nucleus, 2 µm, expanded region, 1 µm. (**B**) STED microscopy images of nuclei and expanded nuclear regions from HEK-293T-FF-EWSR1 (upper) and HT-1080-FF-EWSR1 (lower) cells. Merged images show anti-TAF15 (yellow) and nascent RNA (magenta). Scale bars, nucleus, 2 µm, expanded region,1 µm. (**C, D**) Spatial cluster analysis of the FUS (**C**) and TAF15 (**D**) pattern distributions relative to newly synthesized RNA in the indicated FF-EWSR1 reporter cell lines using EU-labeled RNA as the reference structure. Cumulative frequency plots show analysis of STED microscopy images of four representative nuclei per indicated reporter cell line. (**E, F**) STED microscopy images of nuclei and two expanded regions (i and ii) from HT-1080-FF-EWSR1 cells treated with DMSO (upper panels), or 50 nM (middle panels) or 5000 nM (lower panels) dTAG-13 for 0.5 hrs (left) or 2 hrs (right). Merged images show anti-FUS IF (cyan) and nascent RNA (magenta) (**E**) and anti-TAF15 IF (yellow) and nascent RNA (magenta) (**F**). Scale bars, nucleus, 2 µm, expanded regions i, 1 µm, ii, 0.5 µm. (**G, H**) Spatial cluster analysis of FUS (**G**) and TAF15 (**H**) pattern distributions in HT-1080-FF-EWSR1 cells treated with DMSO or dTAG-13 (50 or 5000 nM) for 0.5 or 2 hrs. Frequency plots show analysis of representative STED microscopy images of a single nucleus per condition per time point per IF analysis. (**I, J**) Spatial cluster analysis of FUS and TAF15 pattern distributions. Frequency plots show analysis of STED microscopy images of four representative nuclei from HT-1080-FF-EWSR1 cels treated with DMSO or dTAG-13 (5000 nM) for 8 hrs (**I**) or 24 hrs (**J**). (**K**) Spatial cluster analysis of FUS and TAF15 pattern distribution in HT-1080-FF-EWSR1 cells using EU-labeled RNA as the reference structure. Cumulative frequency plots show analysis of STED microscopy images of four representative nuclei per condition. (**L**) STED microscopy images of expanded nuclear regions from HT-1080-FF-EWSR1 cells treated with DMSO or dTAG-13 for 24 hrs. Left panels show single channel anti-FUS (cyan) and merged anti-FUS (cyan) and nascent RNA (magenta), right panels show single channel anti-TAF15 (yellow) and merged anti-TAF15 (yellow) and nascent RNA (magenta). Single arrowheads indicate high-intensity FUS or TAF15 signals colocalized with RNA, single arrowed lines indicate lower-intensity FUS or TAF15 signals in proximity to RNA. Scale bars, 1 µm. (**A**, **B, E, F, L**) Images are representative of >20 nuclei per indicated FF-EWSR1 reporter cell line. (**C**,**D**, **G-K**) The dashed line represents a random point pattern distribution; arrowed solid lines indicate deviations from a random point pattern distribution; arrowed dotted lines indicate non-clustered signals (**C**,**D**).

We next evaluated whether EWSR1 depletion promotes the reorganization of FUS and/or TAF15 in a manner consistent with functional compensation. First, we examined FUS and TAF15 using HT-1080-FF-EWSR1 cells treated with DMSO or dTAG-13 (50 or 5000 nM) for 0.5 or 2 hrs. Concomitant with the decrease in the EWSR1 and nucleoplasmic nascent RNA fluorescence (**Figures 6E, 6F** and **S9B, S9C**), representative images of FUS and TAF15 show striking changes in their IF signals (**Figures 6E, 6F**). These results suggest that depletion of EWSR1 induces alterations in the nuclear organization of FUS and TAF15. To quantify changes in the organization of the EWSR1 homologs following depletion of EWSR1, we plotted FUS and TAF15 frequency plots 0.5 and 2 hrs following addition of dTAG-13 (50 and 5000 nM) (**Figure 6G, 6H**). We detected no statistical change in the FUS and TAF15 point pattern distributions following dTAG-13 treatment for 30 min; but after 2 hrs both proteins exhibited a leftward shift in their frequency curves, consistent with increased clustering. We observed similar reorganization of FUS and TAF15 in the TC-32-FF-EWSR1 cells treated with dTAG-13 (**Figures S9D-H**).

Analysis of HT-1080-FF-EWSR1 cells treated with dTAG-13 (5000 nM) for 8 hrs (**Fig. 6I; Fig. S9I**) or 24 hrs (**Figures 6J** and **S9J**) showed progressively enhanced clustering of FUS and TAF15 following EWSR1 degradation. Critically, by 24 hrs, the point pattern distributions of FUS and TAF15 (**Figure 6K**) more closely resembled those of EWSR1 under unperturbed conditions (**Figure 1H**) than those of FUS and TAF15 prior to EWSR1 loss (**Figures 5F** and **5G**).

To determine whether the reorganization of FUS and TAF15 reflected increased association with newly synthesized RNA, we compared FUS and TAF15 clustering to nucleoplasmic nascent RNA point pattern reference structures 24 hrs post addition of dTAG-13, a time point at which nascent RNA levels had returned to pre-treatment levels (**Figure S9J**). In contrast to the partial association with RNA detected in the presence of EWSR1 (**Figures 6C** and **6D**), the point pattern distribution of FUS and TAF15 relative to nucleoplasmic nascent RNA show that upon EWSR1 depletion, FUS and TAF15 exhibited near-complete association with nascent RNA (**Figure 6K**). These cumulative frequency analyses closely matched those observed for baseline EWSR1–RNA interactions (**Figures 2I** and **S4F**), indicating that loss of EWSR1 enables FUS and TAF15 to increase their engagement with nucleoplasmic nascent RNA. Representative images of HT-1080-FF-EWSR1 cells treated with DMSO or dTAG-13 for 24 hrs, shown in **Figure 6L**, highlight the changes in FUS and TAF15 following EWSR1 depletion, including altered detection of both proteins in a distributed state (double arrowheads), as foci (single arrowheads), and an overall increase in their IF signals.

### FUS and TAF15 foci exhibited enhanced colocalization with pS2-RNA pol following EWSR1 degradation

Quantification of FUS and TAF15 nuclear IF intensities, 0.5, 2, 8 and 24 hrs post-addition of dTAG13 showed an increase in the overall nuclear intensity of the IF signals corresponding to both proteins (**Figures 7A** and **S10A**). We therefore asked whether this increase reflected elevated expression of FUS and TAF15. To address this question, we first quantified mRNA levels of *EWSR1*, *FUS*, and *TAF15* at several time points (0.5–24 hrs) following treatment with dTAG-13 (**Figure S10B**). In line with the global suppression of RNA synthesis observed following the initial degradation of EWSR1, transcript levels of all three genes exhibited a reduction at 0.5 hrs, despite the degron system selectively targeting the EWSR1 protein. At 2 and 4 hrs post-treatment, mRNA levels of *EWSR1*, *FUS*, and *TAF15* were comparable to control conditions. Notably, only at 8 and 24 hrs post initiation of dTAG-13 treatment did *FUS* and *TAF15* mRNA levels increase, while *EWSR1* mRNA levels remained unchanged as expected. Immunoblot analysis of nuclear fractionated lysates also detected slight increases in FUS or TAF15 protein abundance after 24 hrs (**Figure S10C**). These data suggest that depletion of EWSR1 modestly enhances expression of FUS and TAF15, but only after several hours.

**Figure 7:**
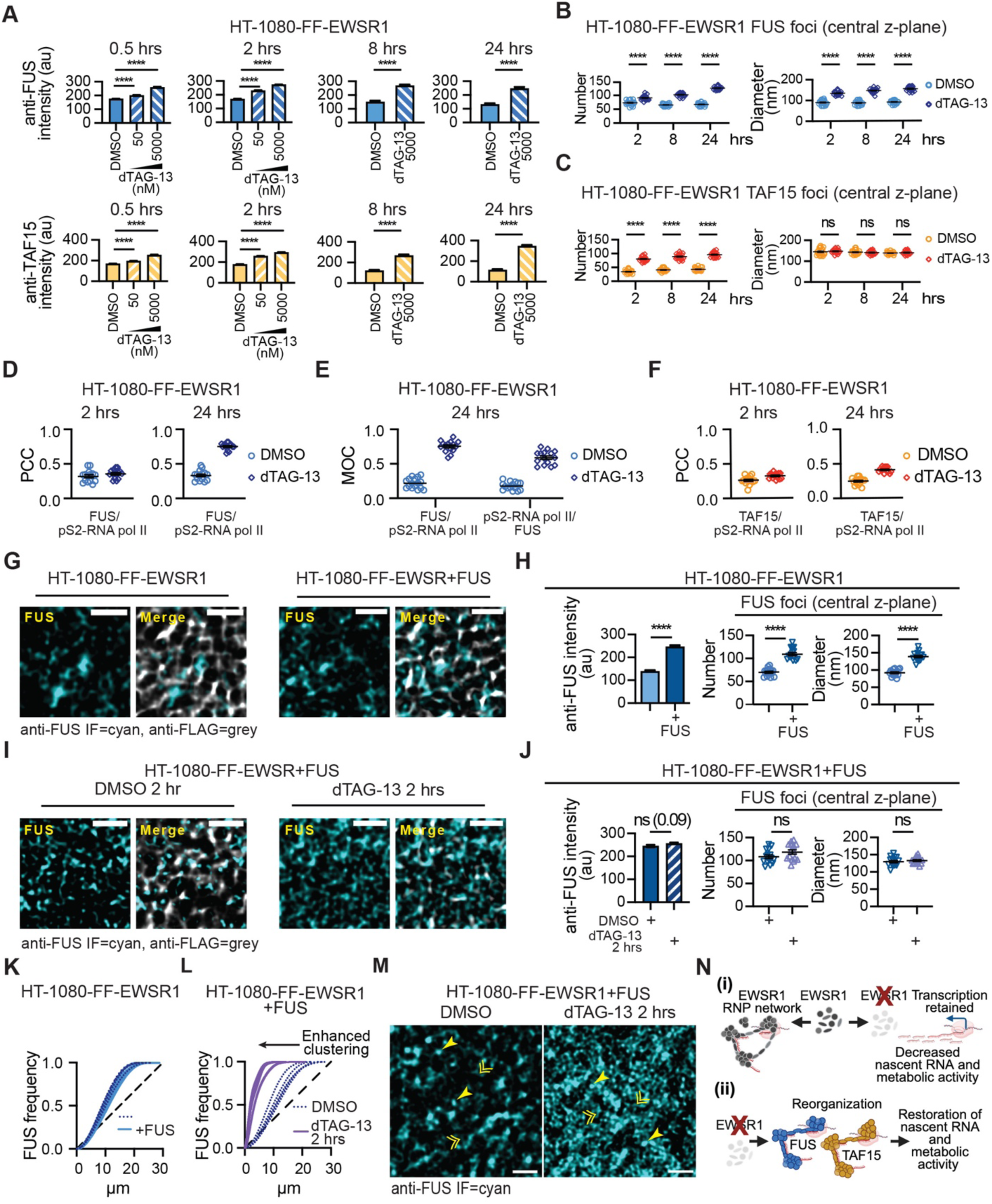
The nuclear reorganization of FUS requires loss of EWSR1. (**A**) Quantification of anti-FUS IF intensity (upper graphs) and anti-TAF15 IF intensity (lower graphs) in HT-1080-FF-EWSR1 cells treated with DMSO or dTAG-13 (as indicated) for the times indicated. (**B, C**) Quantification of the number and diameter (nm) of FUS (**B**) and TAF15 (**C**) foci at the central z-plane of HT-1080-FF-EWSR1 cells treated with DMSO or dTAG-13 (5000 nM) for 2, 8 or 24 hrs. Foci defined as fluorescence intensity ≥5x background. (**D, E**) Quantification of FUS colocalization with pS2-RNA pol II in HT-1080-FF-EWSR1 cells by PCC (**D**) and MOC (**E**) analysis. **(F)** Quantification of TAF15 colocalization with pS2-RNA pol II in TC-32-FF-EWSR1 cells by PCC analysis. **(G)** STED microscopy images (single channel anti-FUS (cyan) and merged (anti-FUS, cyan; anti-FLAG, grey) of an expanded nuclear region from HT-1080-FF-EWSR1 cells and HT-1080-FF-EWSR1 cells stably expressing FUS. Corresponding nuclei images are shown in **Figure S10G**. **(H)** Quantification of anti-FUS intensity, and the number and diameter of FUS foci in HT-1080-FF-EWSR1 cells and HT-1080-FF-EWSR1 cells stably expressing FUS. **(I)** STED microscopy images (single channel anti-FUS (cyan) and merged (anti-FUS, cyan; anti-FLAG, grey) of an expanded nuclear region from HT-1080-FF-EWSR1 cells stably expressing FUS and treated with DMSO or dTAG-13 (5000 nM) for 2 hrs. Corresponding nuclei are shown in **Figure S10H**. **(J)** Quantification of anti-FUS intensity, and the number and diameter of FUS foci in HT-1080-FF-EWSR1 cells stably expressing FUS treated with DMSO or dTAG-13 (5000 nM) for 2 hrs. (**K, L**) Spatial cluster analysis of FUS pattern distribution. Frequency plots show analysis of STED microscopy image of four nuclei from HT-1080-FF-EWSR1 cells and four nuclei from HT-1080-FF-EWSR1 cells stably expressing FUS (**K**) and four nuclei from HT-1080-FF-EWSR1-FUS cells, per condition (DMSO or dTAG-13, 5000 nM for 2 hrs)(**L**). The dashed line indicates a random point pattern distribution, and the arrowed solid lines indicate deviations from a random point pattern distribution. (**M**) STED microscopy images (single channel anti-FUS, cyan) and merged (anti-FUS, cyan; anti-FLAG, grey) of an expanded nuclear region from HT-1080-FF-EWSR1 cells stably expressing FUS and treated with DMSO or dTAG-13 (5000 nM) for 2 hrs. Corresponding nuclei images are shown in **Figure S10K**. (**N**) Schematic of the EWSR1 RNP network required for nascent RNA accumulation and metabolic activity (i) and their restoration of these processes following the reorganization of FUS and TAF15 to form new RNPs (ii). (**A**) Data are shown as mean ± SEM, n=15 nuclei. (**B, C**) Data are shown as individual values and mean ± SEM (black lines), n=20 nuclei per condition per time point. (**D-F**) Data are shown as individual values and mean ± SEM (black lines), n=15 nuclei per treatment per time point. (**H-J**) Intensity data are shown as mean ± SEM and foci number and diameter as individual values and mean ± SEM (black lines), n=15 nuclei per treatment per time point. (**G, I, M**) Images are representative of >10 nuclei per HT-1080-FF-EWSR1 or HT-1080-FF-EWSR1+FUS cell line. Statistical significance was determined using one-way ANOVA (**A,B, C**) and Welch’s t test (**H, J**). (**A, B, C, H, J**) _*_ p<0.05, _***_ p<0.00, _****_ p<0.0001, ns non-significant. Schematic in **(N)** generated using BioRender.

In addition to the potential effects of subtle alterations in protein abundance, we hypothesized that changes in the subnuclear architecture of FUS and TAF15, specifically their redistribution into more organized RNP networks that include concentration at sites of active RNA synthesis, will contribute to the detected increase in fluorescence intensity. Using a ≥5-fold fluorescence intensity threshold to define foci, we quantified FUS and TAF15 foci number and size across 20 nuclei per cell line per condition (DMSO or dTAG-13) at 2, 8, and 24 hours (**Figure 7B** and **7C**). Depletion of EWSR1 resulted in significant increases in both the number and size of FUS foci (**Figure 7B**). Interestingly, following addition of dTAG-13 for 24 hrs, the mean diameter of FUS foci at the central z-plane in HT-1080 cells (156.7 nm) more closely resembled that of the size of EWSR1 foci under unperturbed conditions (mean, 169 nm). In contrast, although the number of TAF15 foci increased following dTAG-13 treatment (**Figure 7C**, left), their diameter remained unchanged (**Figure 7C**, right).

We next examined the relationship between FUS and TAF15 foci and transcriptionally elongating RNA pol II (pS2–RNA pol II). Quantitative colocalization analysis revealed that EWSR1 depletion for 24 hrs led to enhanced colocalization of both FUS and TAF15 foci with pS2–RNA pol II, with a more pronounced effect observed for FUS (**Figures 7D–F** and **S10D-E**). Together, these data indicate that loss of EWSR1 promotes a redistribution of FUS and TAF15 into transcription-associated nuclear foci.

Having further defined the organization of FUS following the degradation of EWSR1, we next sought to address whether such changes depend on protein abundance or required the loss of EWSR1.

### Nuclear reorganization of FUS requires loss of EWSR1

The propensity of FUS to engage in multi-valent interactions is potentially concentration dependent. To determine whether increased FUS protein levels alone can promote the nuclear reorganization of endogenous FUS, or if a reduction in EWSR1 protein levels is needed, we overexpressed FUS in HT-1080-FF-EWSR1 cells (HT-1080-FF-EWSR1+FUS; **Figures 7G**, **S10F,** and **S10G**). FUS overexpression resulted in a 1.75-fold increase in anti-FUS IF intensity relative to control cell, and an increase in both the number and size of FUS foci (**Figure 7H**). Acute degradation of EWSR1 in HT-1080-FF-EWSR1+FUS cells following treatment with dTAG-13 for 2 hrs (**Figures 7I** and **S10H**) did not further increase overall FUS intensity or the number or size of FUS foci (**Figure 7J**). Importantly, while FUS overexpression alone did not alter the point pattern distribution of FUS (**Figure 7K**), depletion of EWSR1 in the HT-1080-FF-EWSR1+FUS cells (**Figures S10I, J**) resulted in a pronounced increase in FUS clustering (**Figure 7L**), suggesting that it is the loss of EWSR1 that is the prime driver of changes in the organization of FUS not protein abundance. Representative images shown in **Figures 7M** and **S10K** illustrate this dramatic change in the organization of FUS stimulated by the loss of EWSR1.

Collectively, these results suggest that under normal physiological conditions, FUS and TAF15 do not compete with EWSR1 for interactions with newly synthesized RNA or its assembly as foci at sites of active transcription. However, upon EWSR1 depletion, FUS and TAF15 undergo spatial reorganization and can functionally compensate by engaging transcription-associated RNA, thereby recapitulating a fundamental function of EWSR1 (**Figure 7N**).

## DISCUSSION

This study demonstrates that formation of an EWSR1-centered RNP network is critical for maintaining nascent RNA levels and reveals that acute loss of EWSR1 activates a compensatory mechanism mediated by its homologs, FUS and TAF15. Both qualitative and quantitative analyses of EWSR1 organization, together with the rapid effects of acute EWSR1 depletion on nascent RNA, support the first component of our working model (**Figure 7N**(i)). Under unperturbed conditions, EWSR1 localizes to discrete nuclear foci that colocalize with sites of active transcription, where it forms the nodes or hubs of an RNP network (**Figures 1**, **2**, and **4**). Acute depletion of EWSR1 underscores the fundamental contributions of this EWSR1-RNP network to the regulation of nascent RNA levels, as it results in a rapid reduction in detectable EU-incorporated RNA accompanied by decreased ATP metabolic activity (**Figure 3**). Notably, a proportional reduction in transcriptional activity, as assessed using a robust marker of RNA pol II elongation does not parallel this loss of nascent RNA (**Figure 4**). These observations suggest that EWSR1 depletion does not overtly affect transcription, but instead, we hypothesize EWSR1 contributes to the stabilization, maturation, or local retention of newly synthesized RNA, potentially through a scaffolding function within the RNP network. Changes in transcriptional output similarly fail to explain the recovery of nascent RNA levels and metabolic activity observed following prolonged (>4 hrs) EWSR1 depletion (**Figures 3** and **4**). Instead, we find that upon depletion of EWSR1, one of its homologs FUS or TAF15 can mediate restoration of metabolic activity (**Figure 5**). This suggests that FUS and TAF15 can compensate for loss of EWSR1, and consistent with this hypothesis, both proteins undergoing pronounced spatial reorganization to assemble RNP networks that resemble the EWSR1-RNP network observed under unperturbed conditions (**Figures 6** and **7**). Accordingly, the second component of our working model (**Figure 7N** (ii)) proposes that compensatory FUS- and TAF15-RNP networks restore nascent RNA levels and mitigate the cellular consequences of EWSR1 depletion and disruption of the endogenous EWSR1-RNP network. Collectively, these findings reveal important distinctions among the physiological roles of FET family proteins while providing the first direct evidence that they can exhibit functional redundancy under specific cellular conditions.

Prior studies of FET proteins, particularly their LCDs, have provided key insights into the multivalent interactions that drive the dynamic concentration of macromolecules into membrane-less compartments, often described as biomolecular condensates.^6,40–45^ Complementary *in vitro* work has further characterized intra- and intermolecular interactions, both in the presence and absence of nucleic acids, that promote self-association of FET proteins into fibril-like assemblies or their incorporation into higher-order nucleic acid–protein complexes.^22,40,41,46,47^ These studies have implicated FET proteins in diverse cellular processes, including transcriptional regulation via interactions with RNA pol II^3,20,21,46,48–50^, RNA processing^51–53^, stress responses and DNA damage and repair, with the latter potentially due to their interactions with RNA:DNA hybrids or R-loops.^25,54–56^ Despite these advances, direct *in vivo* evidence linking the biochemical and biophysical properties of FET proteins to their spatial organization and functional contributions within intact cells has remained limited.^57,58^ The high degree of sequence homology among FET family members, their overlapping nuclear localization patterns, and the difficulty of defining specific RNA interaction parameters, such as sequence, structure, or transcript class, have complicated efforts to delineate their distinct physiological roles.^18,19,59^ Moreover, extensive interactions with other LCD-containing proteins further obscure questions of functional specificity in cellular contexts. By examining a single FET family member at endogenous expression levels, with nanoscale spatial resolution and without overt perturbation of the cellular environment, our study provides *in vivo* support for several aspects of these prior models while refining their relevance to RNA biogenesis and nuclear organization.

Using high-resolution microscopy, we show that EWSR1 consistently forms discrete nuclear foci with minimal variation in number and size across cell lines (**Figures 1E–G** and **Figures S1I**, **S2G**, and **S4C**), although EWS cell lines exhibit larger foci than non-EWS cells. Under unperturbed conditions, FUS and TAF15 similarly display minimal variation in foci number and size (**Figures S7G, S7H**), but following EWSR1 depletion, which reduces the number and size of EWSR1 foci (**Figure S5D**), both FUS and TAF15 show an increase in foci number, while only FUS exhibits a concomitant increase in foci size (**Figure 7B**, **7C**). These observations are consistent with a model in which FET proteins assemble into self-limiting biomolecular condensates governed by reproducible organizational principles rather than stochastic phase separation. Currently, our model only considers the possibility that the FUS, EWSR1, and TAF15 foci are homotypic as depicted in our model (**Figure 7N**), and it will be important in future studies to address whether this is correct, or whether the FET proteins can form heterotypic assemblies within cells.^60^

Another avenue of investigation emerging from our study concerns the structural basis underlying the differential clustering of EWSR1 with nucleoplasmic RNA, compared with FUS and TAF15, under unperturbed conditions, and how this distinction relates to the reorganization of FUS and TAF15 observed upon loss of EWSR1. Previous studies have examined FET protein interactions with RNA and/or DNA in a variety of structural contexts, including in a linear state, as stem–loop and G-quadruplex structures, as well as RNA:DNA hybrids^19,54–56,61–63^. However, translating these *in vitro* observations to the conformation of RNA within cells remains a challenge. As increasingly selective approaches for detecting specific RNA conformational states *in vivo* become available, applying these tools to the model systems used in this study may help define the key features of nascent RNA that preferentially engage one FET protein over another, and how these change following the removal of one of the family members.

It was technically challenging to assign quantitative organizational parameters to the distributed EWSR1 signal frequently observed adjacent to discrete EWSR1 foci (**Figures 1G, 2G**, and **6E–F**), because these signals could not be statistically isolated to evaluate their spatial clustering with nascent RNA. Nonetheless, the presence of this EWSR1 population suggests the coexistence of multiple organizational states within the nucleus. One possibility is that the distributed signal reflects a more elongated or fibrillar mode of assembly, distinct from the compact, focus-like structures analyzed here. This interpretation is consistent with prior electron microscopy and NMR studies demonstrating that FET family proteins can transition between dynamic, liquid-like condensates and more ordered, polymeric or fibrillar assemblies.^40,42,47^ Interestingly, although our quantitative analyses focused on the effects of depletion on EWSR1, FUS, and TAF15 foci (**Figures 7B, 7C**), we also saw a concomitant change in their distributed, non-focal signal populations (**Figures 6L, 7M**). This observation is consistent with a redistribution of these proteins across organizational states and suggests that perturbations affecting condensate formation may also alter the balance between compact and extended assemblies.

The colocalization of EWSR1 foci with pS2–RNA pol II (**Figures 4C–E**), together with the enhanced colocalization of FUS and, to a lesser extent, TAF15 with pS2–RNA pol II following EWSR1 depletion, supports previous proposals that FET proteins function at transcription-associated sites. Nevertheless, despite these spatial relationships, we observed no overt changes in this robust marker of transcriptional elongation following either acute or prolonged EWSR1 depletion (**Figures 4F–I** and **S6C**–**F**). These observations support a model in which recruitment of FET proteins to transcriptionally active regions primarily facilitate interactions with nascent RNA rather than directly modulating transcription itself, although we cannot exclude more subtle contributions to transcriptional regulation. In this context, EWSR1 may function as a molecular scaffold that associates with newly synthesized RNA, thereby supporting RNA stability and promoting the recruitment of additional factors involved in the earliest steps of RNA maturation, which may account for the diversity of functions ascribed to EWSR1. Such a model is consistent with previous descriptions of the *in vitro* analysis of RNA and RBPs interactions seeding the formation of RNA-protein assemblies or condensates.^22,43,64^

Our findings have important implications for diseases associated with FET proteins. Brain tissue from individuals harboring mutations in *FUS* and *TAF15*, and less frequently in *EWSR1*, show evidence of mislocalized FET protein that is aggregated.^9,65,66^ These observations have led to prevailing models in which FET protein aggregation produces a toxic gain-of-function effect that contributes to the phenotype of neurodegenerative disorders such as ALS and FTD. However, it has remained unclear why such mutations do not consistently produce overt loss-of-function phenotypes. Our data suggest that functional redundancy within the FET protein family may buffer against the disrupted function of an individual member. We further speculate that the critical function of the EWSR1-RNP network observed in this study may explain the relative scarcity of disease-associated mutations in *EWSR1* compared with *FUS* and *TAF15*.^66^ Future studies should directly test the compensatory mechanisms identified here in cells harboring disease-associated FET mutations, and address whether EWSR1 can reciprocally substitute for FUS and/or TAF15, or whether FUS and TAF15 can compensate for one another. These questions are also relevant in cancer, where many oncogenic fusion proteins contain the N-terminal domain of a FET protein. For example, studies have suggested that the EWSR1::FLI1 can disrupt wild-type EWSR1 function in EWS cells.^25^ Although we observed no major differences in EWSR1 nucleoplasmic organization between EWS and non-EWS cell lines, aside from increased foci size potentially reflecting enhanced transcriptional activity driven by EWSR1::FLI1 (**Figure S4D**), our experimental systems provide a foundation for future studies examining EWSR1 function in additional contexts including mitosis.^67^ Finally, our findings have implications for therapeutic strategies aimed at targeting FET fusion oncoproteins by disrupting biomolecular condensates.^68,69^ Specifically, our study stresses the need for highly selective targeting that minimizes impact on the wild-type FET proteins as our data indicate that while FUS and TAF15 can compensate for loss of EWSR1, the simultaneous disruption of all the homologous FET proteins is incompatible with the viability of non-transformed as well as tumor cells.

In conclusion, by studying endogenous EWSR1 with quantitative, nanoscale resolution and employing selective, acute degradation to probe temporal and concentration-dependent responses, we define a critical function for an EWSR1-RNP network in the homeostatic maintenance of nascent RNA levels and uncover a compensatory mechanism mediated by FUS and TAF15. These findings provide a unifying framework for understanding FET protein organization, function, and redundancy in both physiological and disease contexts.

### Limitations

Our study describes the nucleoplasmic organization of EWSR1 across multiple cell types using two complementary reporter systems introduced at the endogenous *EWSR1* locus. Although the combined use of CRISPR–Cas9 genome editing and an FKBP12^F36V^ degron system enables analysis of EWSR1 at endogenous protein levels and allows tunable degradation through controlled dTAG-13 treatment, the introduction of any peptide tag into an endogenous protein carries the potential to perturb its native localization, interactions, or function. While our results across two independent tagging strategies support the robustness of our conclusions, we cannot fully exclude subtle tag-induced effects. We also note that in this study, we have utilized the advancements in imaging resolution limits via 2D-STED to assess the spatial relationship of the FET proteins with nascent RNA with a focus on maximizing signals on the central z-plane. Future analysis will use complementary 3D-based imaging to further enhance our understanding of the organization of EWSR1 and its homologs. Also, we conclude that RNA, rather than DNA, is the primary determinant of EWSR1 nucleoplasmic organization. However, our data do not rule out contributions from RNA:DNA hybrids, including R-loops, as an additional organizing factor. Although we did not address this possibility in the current study, the cellular models developed here should provide a framework for investigating this question in the future. Finally, we used pS2-RNA pol II as a marker of global transcriptional activity; however, transcription is a highly complex and dynamic process involving numerous factors and regulatory steps. As such, our analyses provide only a partial snapshot of the transcriptional consequences of EWSR1 depletion. A more comprehensive assessment of how EWSR1 influences global transcriptional programs, including additional stages of transcription and associated regulatory proteins, remains an important goal for future studies.

## RESOURCE AVAILABILITY

### Lead contact

Further information and requests for reagents and resources should be directed to the lead contact, Natasha Caplen

### Materials availability

All reagents and materials are listed in the **key resources table**. All materials generated in this study are available from the lead contact upon request.

### Data and code availability

Unprocessed and uncompressed imaging data have been deposited at Mendeley: 10.17632/wk5tbr8j4f.1. All data deposited will be publicly available as of the date of publication.

This paper does not report any original code

## Supporting information

Resources and Methods

## ACKNOWLEDGEMENTS

We are grateful to members of the Genetics Branch (GB), particularly Ioannis (Yannis) Grammatikakis (Regulatory RNAs and Cancer section, GB) and former and current members of the Functional Genetics Section, GB, particularly Keith Collins and Aaron Kidane, for discussion and comments on the manuscript. We also thank the CCR Genomics Core (Sanger and whole plasmid Oxford Nanopore sequencing) and CCR Flow Cytometry Core for technical assistance. This work was supported by the Intramural Research Program of NCI, NIH to N.J.C, ZIA: BC 011704, and ZIC BC 010858 supports the CCR Confocal Microscopy Core Facility. The generation of CRISPR-Cas9 reagents included the support of Federal funds from the NCI, NIH, under Contract No. HHSN261201500003I.

The content of this publication does not necessarily reflect the views or policies of the US Department of Health and Human Services, nor does mention of trade names, commercial products, or organizations imply endorsement by the US Government.

## AUTHOR CONTRIBUTIONS

Conceptualization : S.S.R and N.J.C; Methodology: S.S.R, A.T, M.J.K, and N.J.C; Investigation: S.S.R, I.K, T.L.J, T.B, V.J.E, Resources: L.L, A.T, M.J.K and N.J.C; Writing original draft: S.S.R and N.J.C; Writing review and editing: all authors; Supervision: N.J.C; Funding acquisition: N.J.C.

## DECLARATION OF GENERATIVE AI AND AI-ASSISTED TECHNOLOGIES

During the preparation of this work, the authors used NIH CHIRP (ChatGPT) to edit the text for clarity and length. After using this tool, the authors reviewed and edited the content as needed and take full responsibility for the content of the publication.

