## Supplementary Figures for "FUS and TAF15 safeguard the critical functions of the ribonucleoprotein network formed by EWSR1 and newly synthesized RNA"

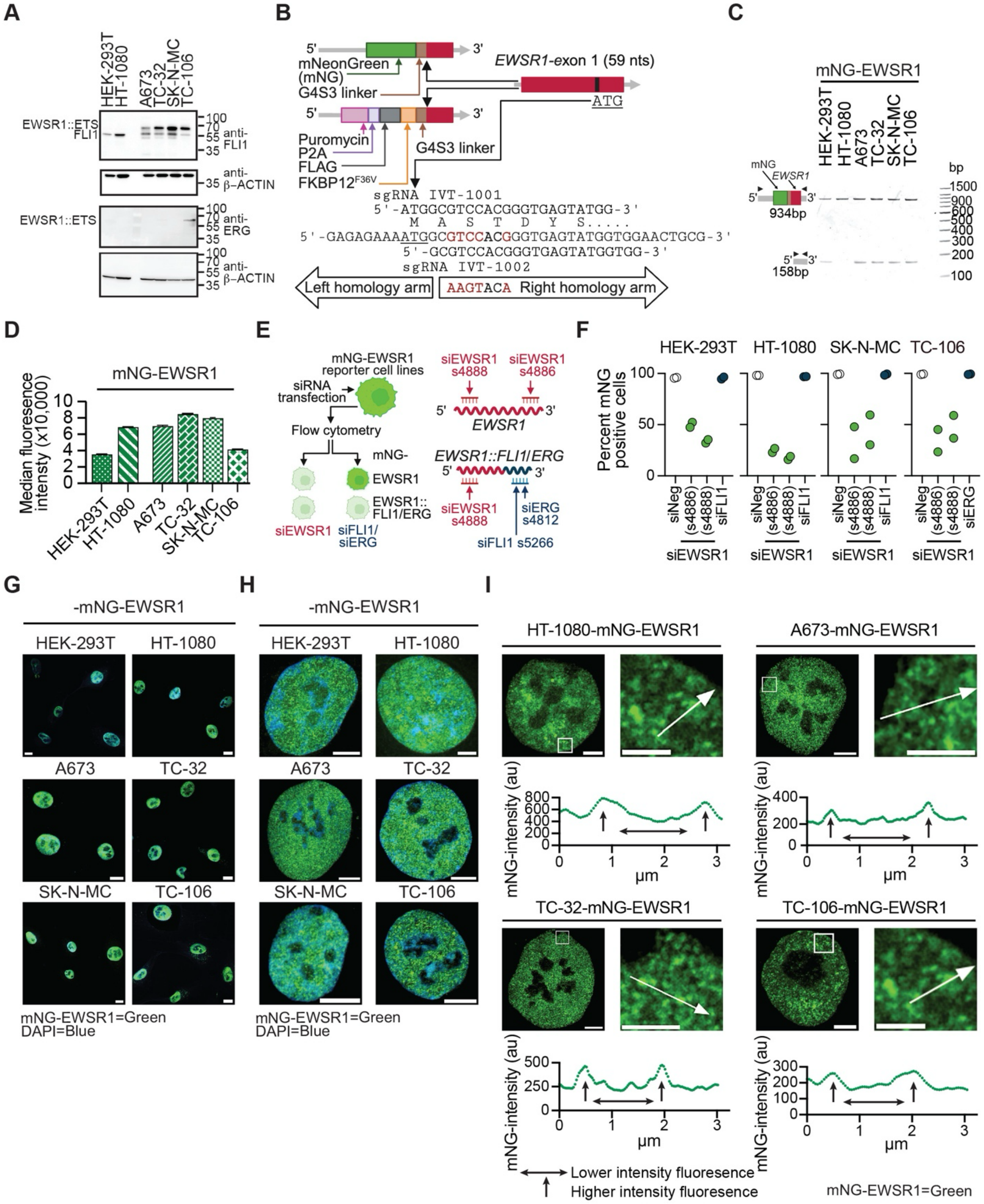


**Figure S1: Generation and validation of mNeonGreen-EWSR1 (mNG-EWSR1) reporter cell lines**

(**A**) Immunoblot analysis of whole cell lysates from the indicated cell lines probed with antibodies against the indicated proteins.

(**B**) Schematic of the CRISPR-Cas9-mediated modification of the endogenous *EWSR1* locus illustrating insertion of indicated DNA cassettes immediately 5' of *EWSR1* exon 1. Red text denotes sequence changes introduced into exon 1 by donor construct.

(**C**) PCR-based analysis confirming integration of the mNG reporter cassette into the *EWSR1* locus. PCR primers amplify across the donor cassette and flanking 5' and 3' homology arms, indicating. monoallelic modification in HEK-293T cells, biallelic modification in HT-1080 cells, and monoallelic modification in EWS cell lines.

(**D**) Flow cytometry analysis showing median mNG fluorescence intensity in the indicated mNG-EWSR1 reporter cell lines.

(**E**) Schematic of the siRNA-based assay used to determine which *EWSR1* alleles are modified, using siRNAs targeting *EWSR1* or *EWSR1::FLI1/ERG* transcripts[1-3].

(**F**) Flow cytometry analysis following siRNA transfection (two independent replicates). In SK-N-MC and TC-106 cells, reduction in mNG positive cells following transfection of siEWSR1 but not siFLI1 or siERG indicates modification of the unarranged *EWSR1* allele. Corresponding analyses in A673-mNG-EWSR1 and TC-32-mNG-EWSR1 reporter cell lines are reported in Rajan *et al.*, 2024[3].

(**G**) Confocal images of nuclei showing mNG–EWSR1 fluorescence (green) and DAPI-stained DNA (blue). Scale bar, 25 μm.

(**H**) SoRa super resolution confocal images of whole showing mNG-EWSR1 fluorescence (green). Scale bar, 4 μm.

(**I**) SoRa super resolution confocal images showing nuclei (left) and expanded nuclear regions (right) with corresponding line plots. mNG-EWSR1 fluorescence (green) and line plots correspond to the white lines shown in the expanded regions (arrow direction indicates the x-axis orientation of each plot). Horizontal arrowed lines indicate lower intensity mNG-EWSR1 signal. Vertical arrowed lines indicate higher intensity signal. Scale bars, nucleus, 4 μm, expanded region, 2 μm.

(**G**-**I**) Images are representative of >20 cells or nuclei per indicated mNG-EWSR1 reporter cell line.

Schematics in (**B**), (**C**), and (**E**) were generated using BioRender.

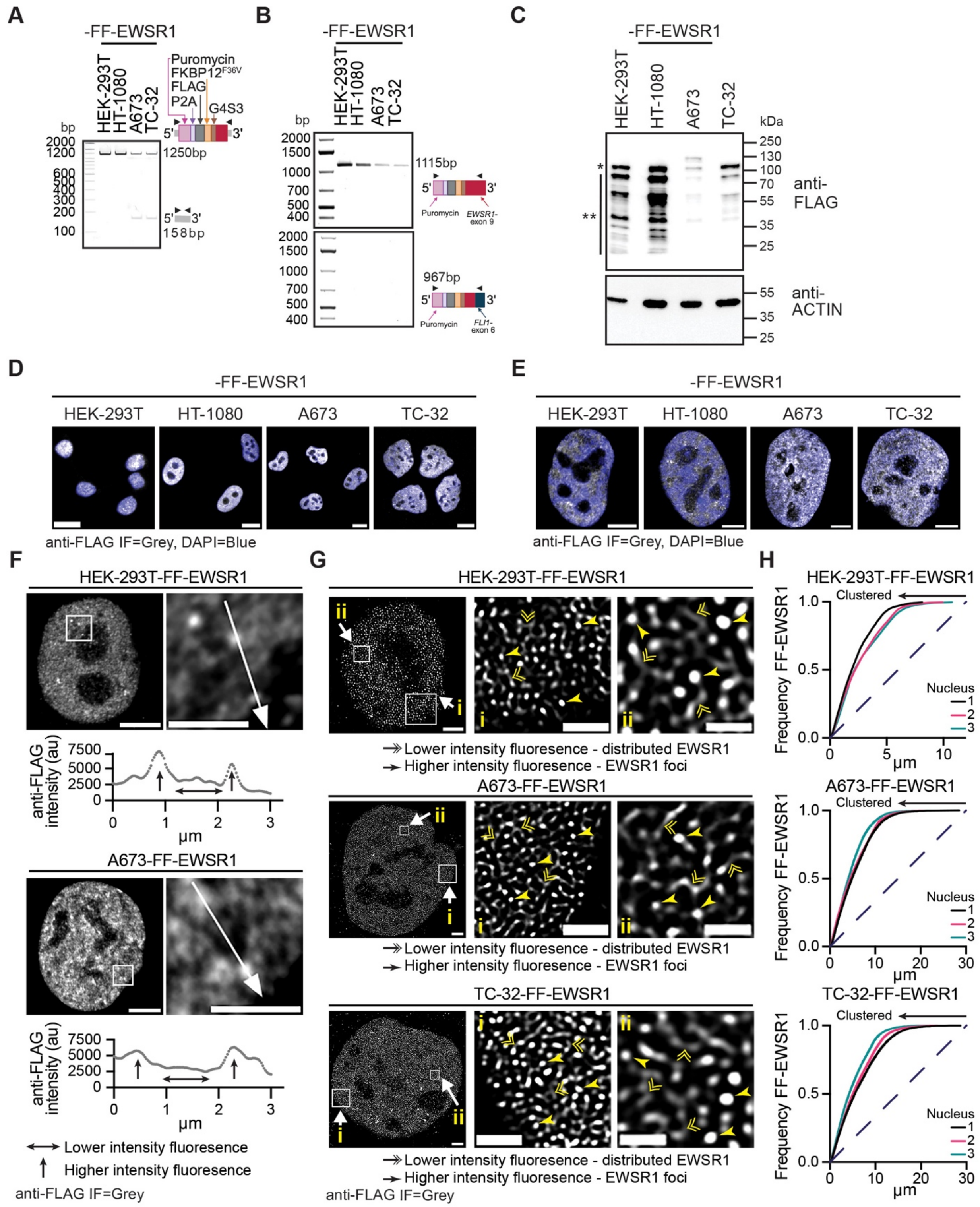


**Figure S2: The generation of FLAG-FKBP12$^{F36V}$-EWSR1 (FF-EWSR1) reporter cell lines**

(**A**) PCR-based analysis confirming targeted integration of the FF cassette into the endogenous *EWSR1* locus in the indicated reporter cell lines. PCR primers were designed to amplify across the donor cassette and the flanking 5' and 3' genomic regions encompassed by the *EWSR1* homology arms. This analysis demonstrates biallelic modification of *EWSR1* in HEK-293T and HT-1080 cells and monoallelic modification in EWS cell lines.

(**B**) RT-PCR analysis of FF-*EWSR1* transcripts using a common puromycin-specific forward primer and reverse *EWSR1*-exon 9 or *FLI1*-exon 6 primers. Modified *EWSR1* transcripts are detected, whereas modified *EWSR1::FLI1* transcripts are not.

(**C**) Immunoblot analysis showing an ~30 kDa molecular weight increase of FF-modified EWSR1. * and ** indicate modified major and minor EWSR1 isoforms, respectively.

(**D**) Confocal images of nuclei from the indicated FF-EWSR1 reporter cell lines showing anti-FLAG (grey) and DAPI (blue). Scale bar, 25 μm.

(**E**) SoRa super resolution confocal images of nuclei from the indicated FF-EWSR1 reporter cell lines showing anti-FLAG (grey) and DAPI (blue). Scale bar, 4 μm.

(**F**) SoRa super resolution confocal images showing nuclei (left) and expanded nuclear regions (right with corresponding line plots. anti-FLAG IF (grey) and line plots correspond to the white lines shown in the expanded regions (arrow direction indicates the x-axis orientation of each plot). Horizontal arrowed lines indicate lower anti-FLAG signal, whereas vertical arrowed lines indicate higher intensity signal. Scale bars, nucleus, 4 μm, expanded region, 2 μm.

(**G**) STED microscopy images of nuclei and expanded nuclear regions (i and ii) from the indicated FF-EWSR1 reporter cell lines showing anti-FLAG IF (grey). Double arrowheads indicate lower-intensity distributed FF-EWSR1 signal and single arrowheads indicate high intensity FF-EWSR1 foci. Scale bars, nucleus, 2 μm, expanded regions i, 1 μm, ii, 0.5 μm.

(**H**) Spatial cluster analysis of FF-EWSR1 pattern distribution in the indicated cell lines. Frequency plots show analysis of STED microscopy images of four nuclei. The dashed line indicates a random point pattern distribution; arrowed solid lines indicate deviations from a random point pattern distribution.

(**D**-**G**) Images are representative of > 20 cells or nuclei per indicated FF-EWSR1 reporter cell line.

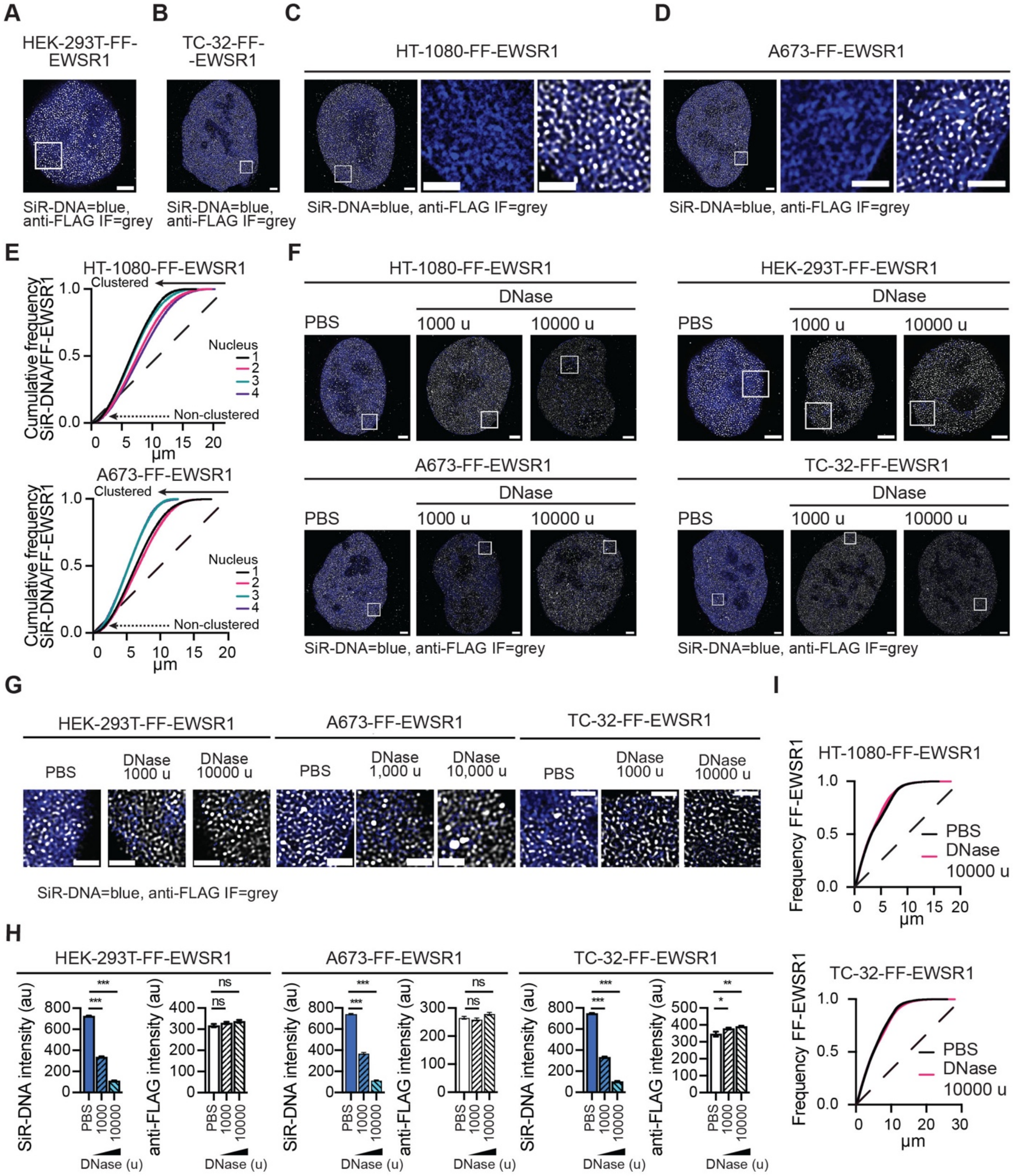


**Figure S3: The nuclear organization of EWSR1 is DNA-independent**

(**A, B**) STED microscopy images of nuclei from HEK-293T–FF–EWSR1 (**A**) and TC-32–FF–EWSR1 cells (**B**) with the boxes indicating the expanded regions shown in **Figures 2A** and **2B**, respectively. Merged images show the FLAG IF (grey) and SiR-DNA (blue). Scale bars, 2 µm.

(**C, D**) STED microscopy images of nuclei and indicated expanded regions from HT-1080–FF–EWSR1 (**C**) and A673–FF–EWSR1 (**D**) cells. Expanded regions show single-channel SiR-DNA (blue) and merged SiR-DNA (blue) with anti-FLAG IF (grey). Scale bars, nucleus, 2 µm, expanded region, 1 µm.

(**E**) Spatial cluster analysis of FF-EWSR1 IF pattern distribution relative to DNA, using SiR-DNA as the reference structure. Cumulative frequency plots show analysis of STED microscopy images of four representative nuclei per indicated reporter cell line.

(**F**) STED microscopy images of nuclei from the indicated FF-EWSR1 reporter cell lines treated with PBS, or DNase (1000 or 10000 u, 20 minutes (min)). Boxes indicate the expanded regions shown in **Figures 2D** or **Figure S3G**. Merged images show anti-FLAG IF (grey) and SiR-DNA (blue). Scale bar, 2 μm.

(**G**) Expanded regions from HEK-293T-FF-EWSR1, A673-FF-EWSR1, and TC-32-FF-EWSR1 cells treated with PBS or DNase (1000 or 10000 u, 20 min). Merged images show anti-FLAG IF (grey) and SiR-DNA (blue). Corresponding whole-nucleus images are shown in **Figure S3F**. Scale bar, 1 μm.

(**H**) Quantification of SiR-DNA and anti-FLAG fluorescence intensities in the indicated FF-EWSR1 reporter cells following treatment with either PBS or DNase (1000 or 10000 u, 20 min).

(**I**) Spatial cluster analysis of FF-EWSR1 IF pattern distribution. Plots show analysis of STED microscopy images of representative nuclei from HT-1080-FF-EWSR1 and TC-32-FF-EWSR1 cells treated with PBS or DNase (10000 u).

(**A**-**D, F, G**) Images are representative of >20 nuclei per indicated FF-EWSR1 reporter cell line.

(**E**, **I**) The dashed line indicates a random point pattern distribution; arrowed solid line indicates deviations from a random point pattern distribution; arrowed dotted lines indicate non-clustered signals (**E**).

(**H**) Data are shown as mean ± SEM from 20 nuclei per treatment per reporter cell line. Statistical significance was determined by one-way ANOVA, ⋆ p<0.05, ⋆⋆ p<0.01, ⋆⋆⋆ p<0.001, ns non-significant.

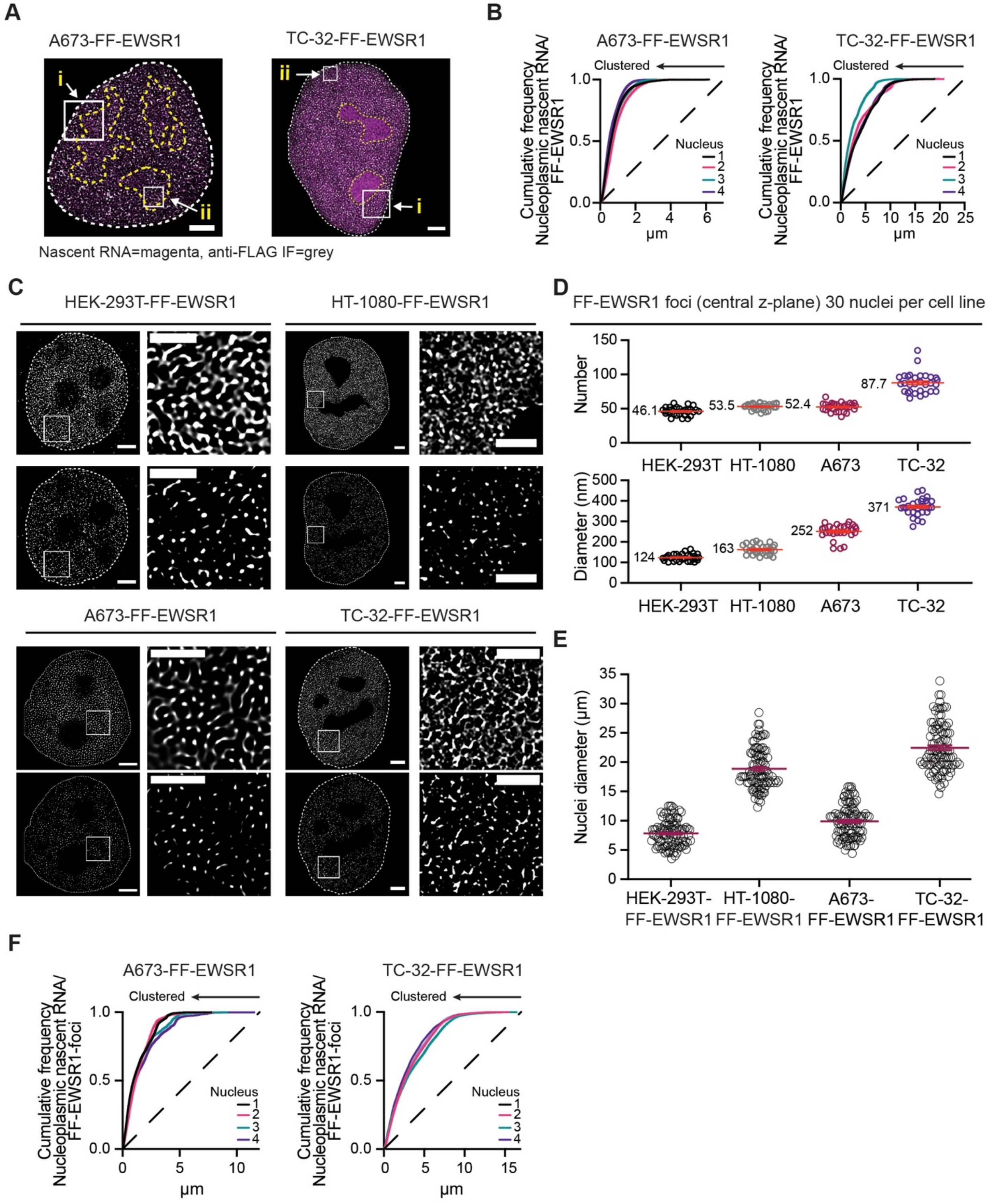


**Figure S4: EWSR1 and newly synthesized RNA form a ribonucleoprotein network**

(**A**) STED microscopy images of nuclei from A673-FF-EWSR1 and TC-32-FF-EWSR1 cells. Merged images show anti-FLAG IF (grey) and nascent RNA labeling (magenta). The white boxes indicate the expanded regions shown in **Figure 2G**. Scale bars, 2 μm.

(**B**) Spatial cluster analysis of FF-EWSR1 IF pattern distribution relative to newly synthesized RNA in the indicated FF-EWSR1 reporter cell lines using EU-labeled RNA as the reference structure.

Cumulative frequency plots show analysis of STED microscopy images of four representative nuclei per indicated reporter cell line.

(**C**) STED microscopy images of nuclei and expanded nuclear regions from the indicated FF-EWSR1 reporter cell lines. Upper panels show anti-FLAG IF signals (grey) above background and lower panels show FF–EWSR1 foci (grey) defined by ≥5× fluorescence intensity relative to background. Scale bars, nucleus, 2 μm, expanded region, 1 μm.

(**D**) Quantification of the number and the diameter (nm) of FF-EWSR1 foci at the central z-plane of the nuclei from the indicated reporter cell lines. Foci defined as anti-FLAG IF signal ≥5x above background. n=30 nuclei per indicated reporter cell line. Numbers indicate the mean for each parameter in each reporter cell line

(**E**) Quantification of the nuclear diameter (nm) in the indicated FF-EWSR1 reporter cell lines. n=100 nuclei per cell line.

(**F**) Spatial cluster analysis of high intensity (≥5x signal) FF-EWSR1 foci pattern distribution relative to newly synthesized RNA in the indicated FF-EWSR1 reporter cell lines using EU-labeled RNA as the reference structure. Cumulative frequency plots show analysis of STED microscopy images of four representative nuclei per indicated reporter cell line.

(**A**, **C**) Images are representative of >20 nuclei per indicated FF-EWSR1 reporter cell line.

(**B**, **F**) The dashed lines indicate a random point pattern distribution; arrowed solid lines indicate deviations from a random point pattern distribution.

(**D**) n=30 nuclei per indicated reporter cell line. Data show as individual data points and mean ±SEM (red lines). Values indicate the mean for each parameter in each cell line.

(**E**) n=100 nuclei per reporter cell line. Data show as individual data points and mean ±SEM (red lines).

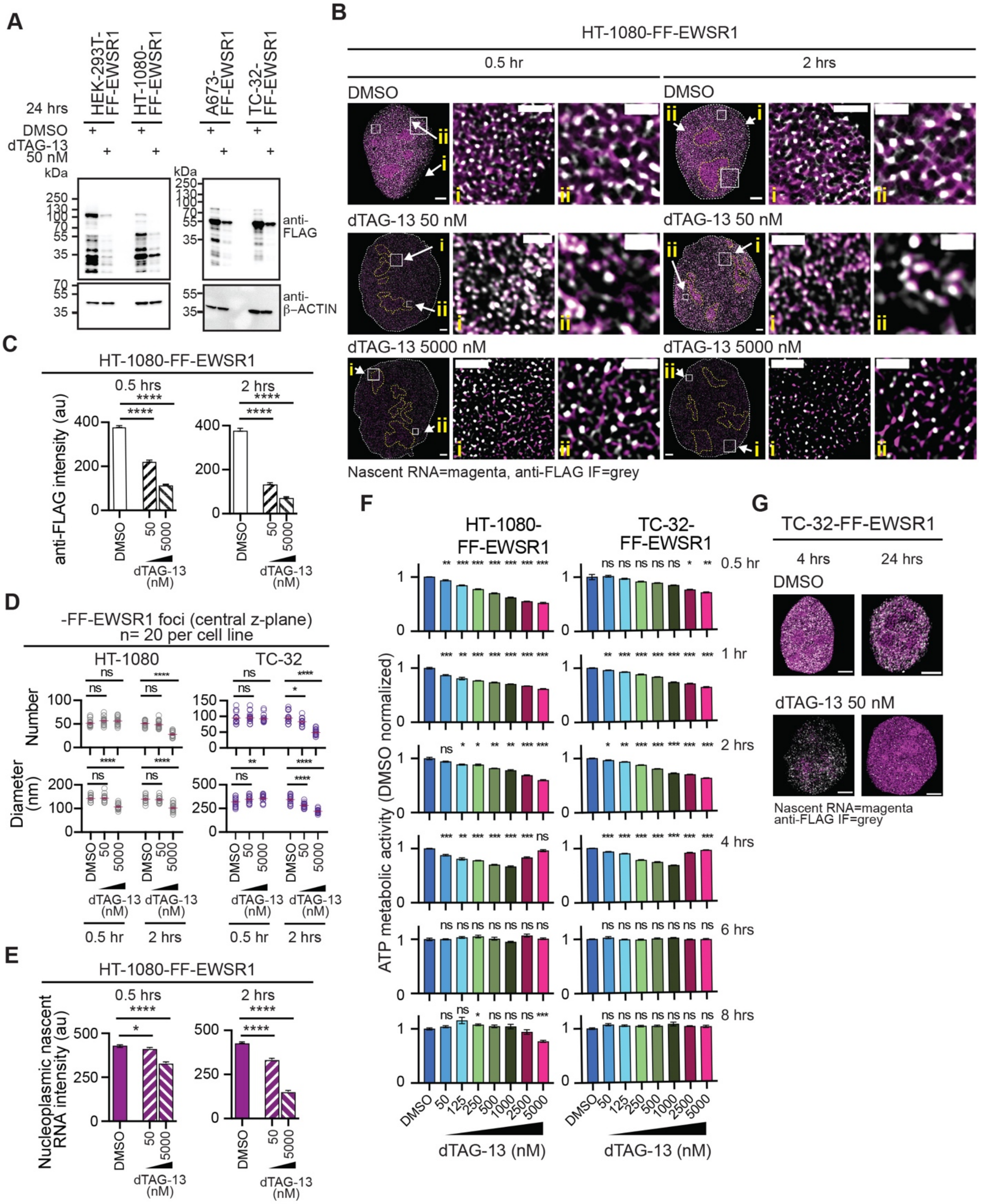


**Figure S5: The degradation of EWSR1 results in a transient decrease in nascent RNA and cell viability.**

(**A**) Immunoblot analysis of whole cell lysates from the indicated FF-EWSR1 reporter cell lines treated with DMSO or dTAG-13 (50 nM) for 24 hours (hrs) and probed with antibodies against the indicated proteins.

(**B**) STED microscopy images of nuclei and two expanded regions (i and ii) from HT-1080-FF-EWSR1 cells treated with DMSO (upper panels), or dTAG-13 (50 nM, middle panels, or 5000 nM, lower panels) for 0.5 or 2 hrs. Merged images show anti-FLAG IF (grey) and nascent RNA (magenta). Scale bars, nucleus 4 μm, expanded regions i, 1 μm, ii, 0.5 μm.

(**C**) Quantification of anti-FLAG IF intensity in HT-1080-FF-EWSR1 cells following treatment with DMSO or dTAG-13 (50 or 5000 nM) for 0.5 (left) or 2 (right) hrs.

(**D**) Quantification of the number and the diameter (nm) of FF-EWSR1 foci measured at the central z-plane of the indicated reporter cell lines following treatment with DMSO or dTAG-13 (50 or 5000 nM) for 0.5 or 2 hrs. Foci were defined as anti-FLAG fluorescence intensity ≥5x above background.

(**E**) Quantification of nascent RNA fluorescence intensity in HT-1080-FF-EWSR1 cells following treatment with DMSO or dTAG-13 (50 or 5000 nM) for 0.5 (left) or 2 (right) hrs.

(**F**) Cell viability of HT-1080-FF-EWSR1 and TC-32-FF-EWSR1 reporter cell lines measured 0.5 – 8 hrs after addition of DMSO or increasing concentration of dTAG-13 (50 -5000 nM). Viability values are normalized to the mean of DMSO-treated cells at each time point.

(**G**) SoRa images of nuclei from TC-32-FF- EWSR1 reporter cells 4 or 24 hrs after treatment with DMSO or dTAG-13 (50 nM). Scale bars, 4 μm. Quantification of anti-FLAG IF and nascent RNA fluorescence intensities is shown in **Figure 3E**.

(**B, G**) Images are representative of >20 nuclei per indicated FF-EWSR1 reporter cell line.

(**C, E**, **F**) Data shown as mean ± SEM. (**D**) Individual data points (circles) and the mean ± SEM (lines) (**C, E**) n=15 nuclei per reporter cell line per condition. (**D**) n=20 nuclei per reporter cell line. (**F**) n=6 biological replicates per reporter cell line per treatment per time point.

(**C-F**) Statistical significance was determined using one-way ANOVA. (**C -F**) * p<0.05, ** p<0.01, *** p<0.00, **** p<0.0001, ns non-significant.

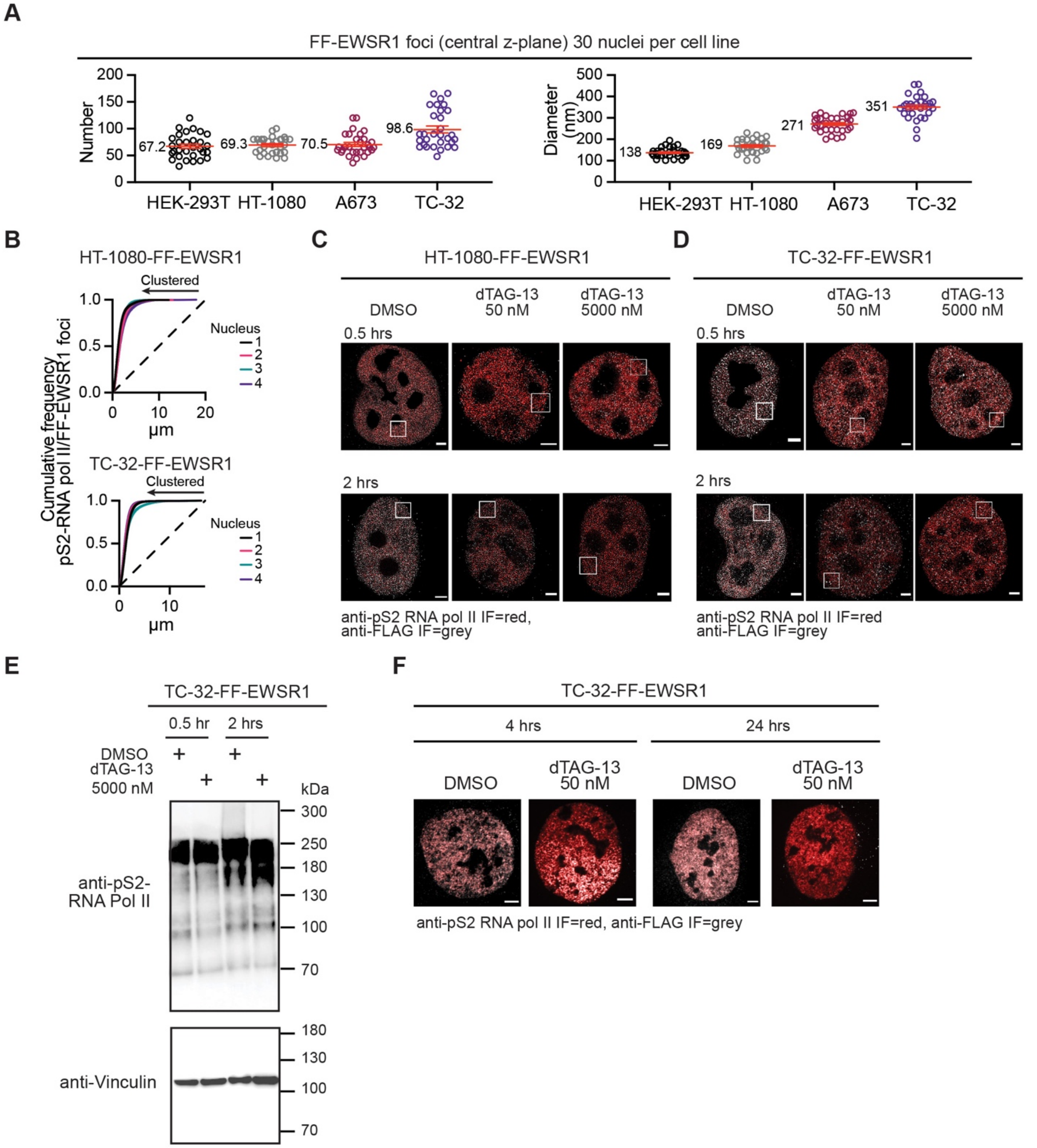


**Figure S6: EWSR1 foci colocalize with actively elongating RNA polymerase II, but depletion of EWSR1 does not measurably alter this marker of transcriptional activity**

(**A**) Quantification of the number and diameter (nm) of FF-EWSR1 foci at the central z-plane of the indicted cells. Foci defined as anti-FLAG fluorescence intensity ≥5x than background.

(**B**) Spatial cluster analysis of FF–EWSR1 foci (≥5x above background) pattern distribution as a function of distance relative to pS2-RNA pol II IF in the indicated FF–EWSR1 reporter cell lines. Cumulative frequency plots were generated from STED microscopy images of four nuclei per reporter cell line.

(**C, D**) STED microscopy images of nuclei from HT-1080-FF-EWSR1 cells (**C**) and TC-32-FF-EWSR1 cells (**D**) treated with DMSO or dTAG-13 (50 or 5000 nM) for the indicated times. Merged images show anti-FLAG (grey) and pS2-RNA pol II IF (red). Boxes indicate the expanded regions shown in **Figure 4F**. Scale bar, 2 µm.

(**E**) Immunoblot analysis of whole cell lysates from TC-32-FF- EWSR1 cells treated with either DMSO or dTAG-13 (5000 nM) for the indicated times and probed using the antibodies against the indicated proteins.

(**F**) SoRa images of the nuclei of the indicated FF- EWSR1 reporter cells 4 or 24 hrs post-addition of DMSO or dTAG-13 (50 nM). Merged images show anti-FLAG (grey) and pS2-RNA pol II IF (red). Scale bar, 4 µm.

(**A**) Data shown as individual data points and mean ± SEM (red lines), mean value indicated. n=30 nuclei per reporter cell line.

(**B**) The dashed line represents a random point pattern distribution. Arrowed solid lines indicate deviations from a random point pattern distribution.

(**C**, **D, F**) Images are representative of >20 nuclei per indicated FF-EWSR1 reporter cell line.

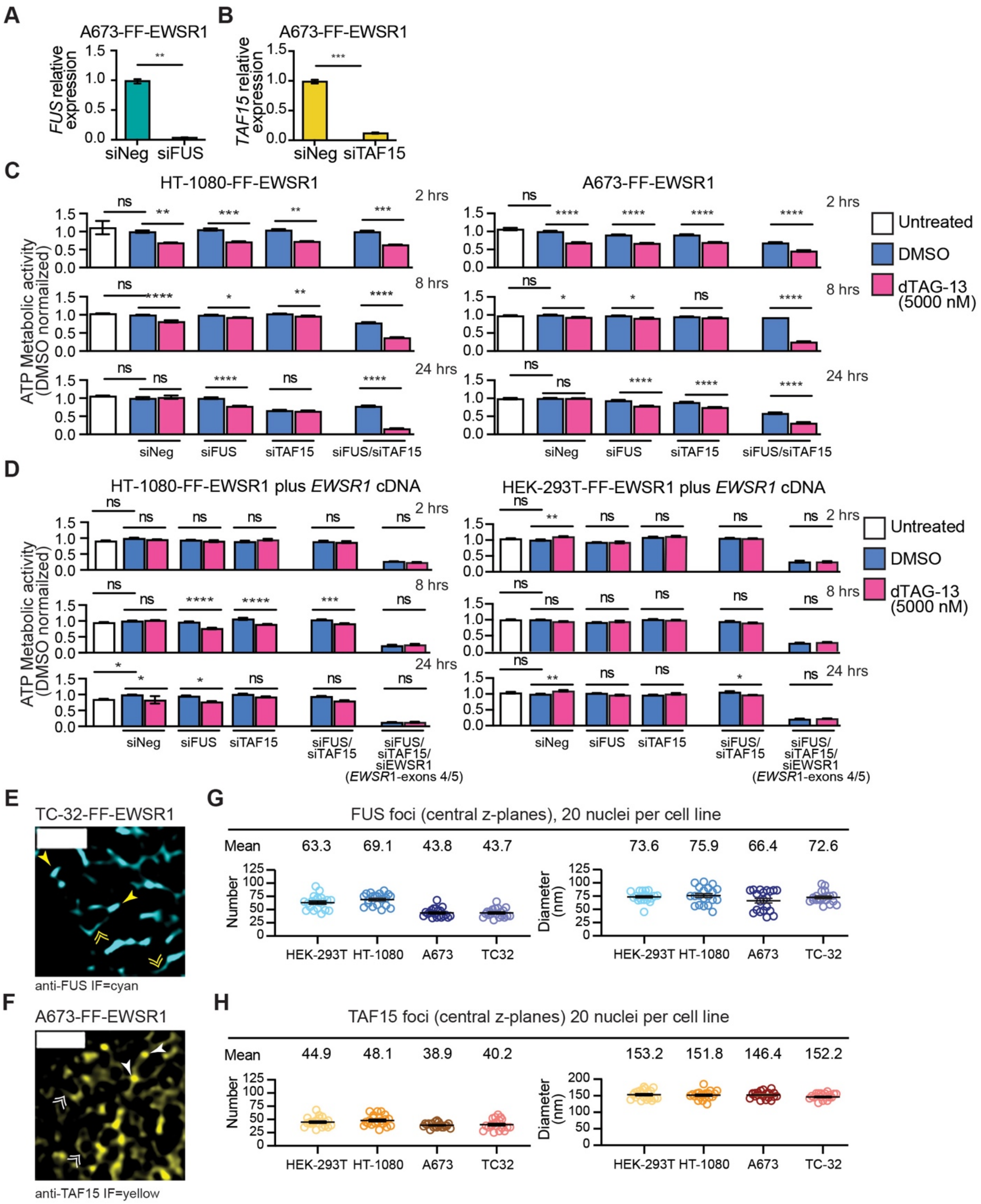


**Figure S7: FUS and TAF15 functionally compensate for EWSR1 loss to restore cellular metabolic activity**

(**A, B**) qRT-PCR analysis of *FUS* (**A**) and *TAF15* (**B**) expression 48 hours post-transfection of A673-FF-EWSR1 cells with the indicated siRNAs. *FUS* or TAF15 expression normalized to the expression of the housekeeping gene *RPL27* and shown relative to siNeg-transfected cells.

**(C)** The ATP metabolic activity of HT-1080-FF-EWSR1 and A673-FF-EWSR1 reporter cells following siRNA-mediated depletion of FUS and/or TAF15 (48 hrs) and dTAG-13-mediated degradation of EWSR1 (2, 8, or 24 hrs). Data are normalized to the mean of the siNeg-transfected, DMSO-treated cells.

(**D**) The ATP metabolic activity of HT-1080-FF-EWSR1 and HEK-293T-FF-EWSR1 reporter cells stably expressing an *EWSR1* cDNA following siRNA-mediated depletion of FUS and/or TAF15 (48 hrs) and dTAG-13-mediated degradation of EWSR1 (2, 8, or 24 hrs). Data are normalized to the mean of the siNeg-transfected, DMSO-treated cells.

(**E**) STED microscopy images of an expanded nuclear region from a TC-32-FF-EWSR1 showing FUS IF (cyan). The double arrowheads indicate lower intensity fluorescence or distributed FUS, and the single arrowheads indicate higher intensity fluorescence or FUS foci. Scale bars, nucleus, 2 μm, expanded region, 1 μm.

(**F**) STED microscopy images of an expanded nuclear region from an A673-FF-EWSR1 showing TAF15 IF (yellow). The double arrowheads indicate lower intensity fluorescence or distributed TAF15, and the single arrowheads indicate higher intensity fluorescence or TAF15 foci. Scale bars, nucleus, 2 μm, expanded region, 1 μm.

(**G, H**) Quantification of FUS (**G**) and TAF15 (**H**) foci number and diameter (≥5× background).

(**A, B**) Data shown as mean ± SEM, n=3 per transfection.

(**C, D**) Data shown as the mean ± SEM, n=6 per treatment.

(**E, F**) Images are representative of >20 nuclei per indicated EWSR1 reporter cell line.

(**G, H**) Data are shown as individual values and mean ± SEM, n=20 nuclei per condition per reporter cell line.

(**A, B**) Statistical significance was determined using Welch's t-test. (**C, D**) Statistical significance was determined using one-way ANOVA.* p<0.05, ** p<0.01, *** p<0.001, **** p<0.0001, ns non-significant.

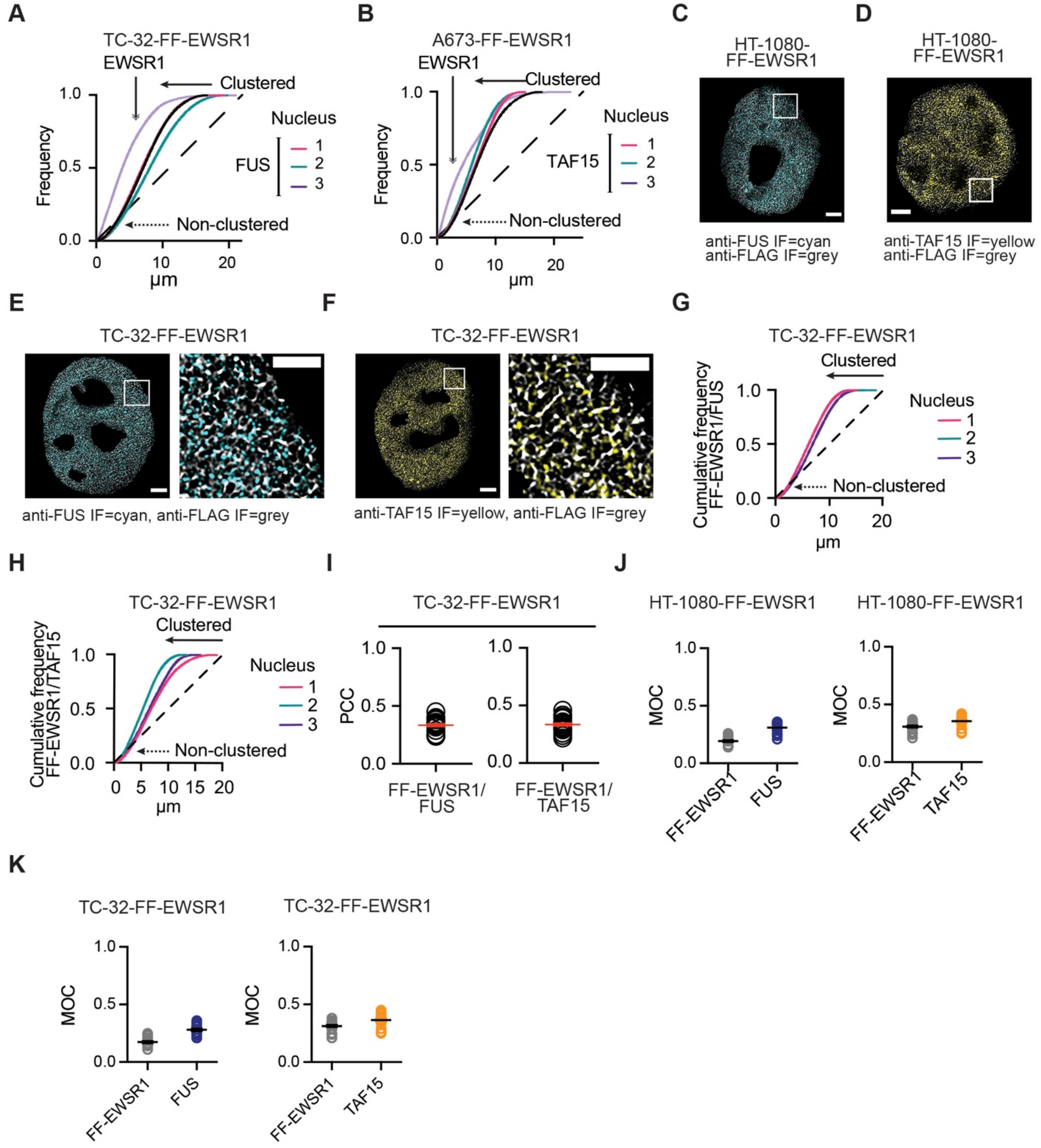


**Figure S8: EWSR1 exhibits minimal spatial colocalization with FUS or TAF15**

(**A, B**) Spatial cluster analysis of FUS IF (**A**) and TAF15 IF (**B**) pattern distributions across distance for TC-32-FF-EWSR1 cells (**A**) and A673-FF-EWSR1 cells (**B**). Frequency plots were generated from STED microscopy images of three nuclei per reporter cell line. A frequency plot of FF-EWSR1 in each report cell line is shown for comparison (light purple).

(**C, D**) STED microscopy images of nuclei from HT-1080-FF-EWSR1 cells. Merged images show anti-FUS (cyan) and anti-FLAG IF (grey)(**C**) or anti-TAF15 (yellow) and anti-FLAG IF (grey) (**D**). Boxes indicate the expanded regions shown in **Figures 5H, I**. Scale bar, 2 µm.

(**E, F**) STED microscopy merged images of nuclei and enlarged nuclear regions from TC-32-FF-EWSR1 cells showing anti-FUS (cyan) and anti-FLAG IF (grey) (**E**) or anti-TAF15 (yellow) and anti-FLAG IF (grey) **(F)**. Scale bars, nucleus, 2 μm, expanded region, 1 μm.

(**G, H**) Spatial cluster analysis of FUS IF (**G**) and TAF15 IF (**H**) pattern distributions as a function of distance relative to FF-EWSR1 in TC-32-FF-EWSR1 cells. Cumulative frequency plots were generated from STED microscopy images of three nuclei per reporter cell line per analysis.

(**I**) Quantification of FF–EWSR1 colocalization with FUS or TAF15 in TC-32-FF-EWSR1 cells by PCC analysis.

(**J**) Quantification of FF–EWSR1 colocalization with FUS or TAF15 in HT-1080-FF-EWSR1 cells by MOC analysis.

(**K**) Quantification of FF–EWSR1 colocalization with FUS or TAF15 in TC-32-FF-EWSR1 cells by MOC analysis.

(**A**, **B**, **G**, **H**) The dashed line represents a random point pattern distribution; arrowed solid lines indicate deviations from a random point pattern distribution; arrowed dotted lines indicate non-clustered signals.

(**C**-**F**) Images are representative of >20 nuclei per indicated EWSR1 reporter cell line.

(**I-K**) Data are shown as individual values with mean ± SEM indicated (**I**, red lines, **J**, **K**, black lines), n=20 nuclei per condition per analysis.

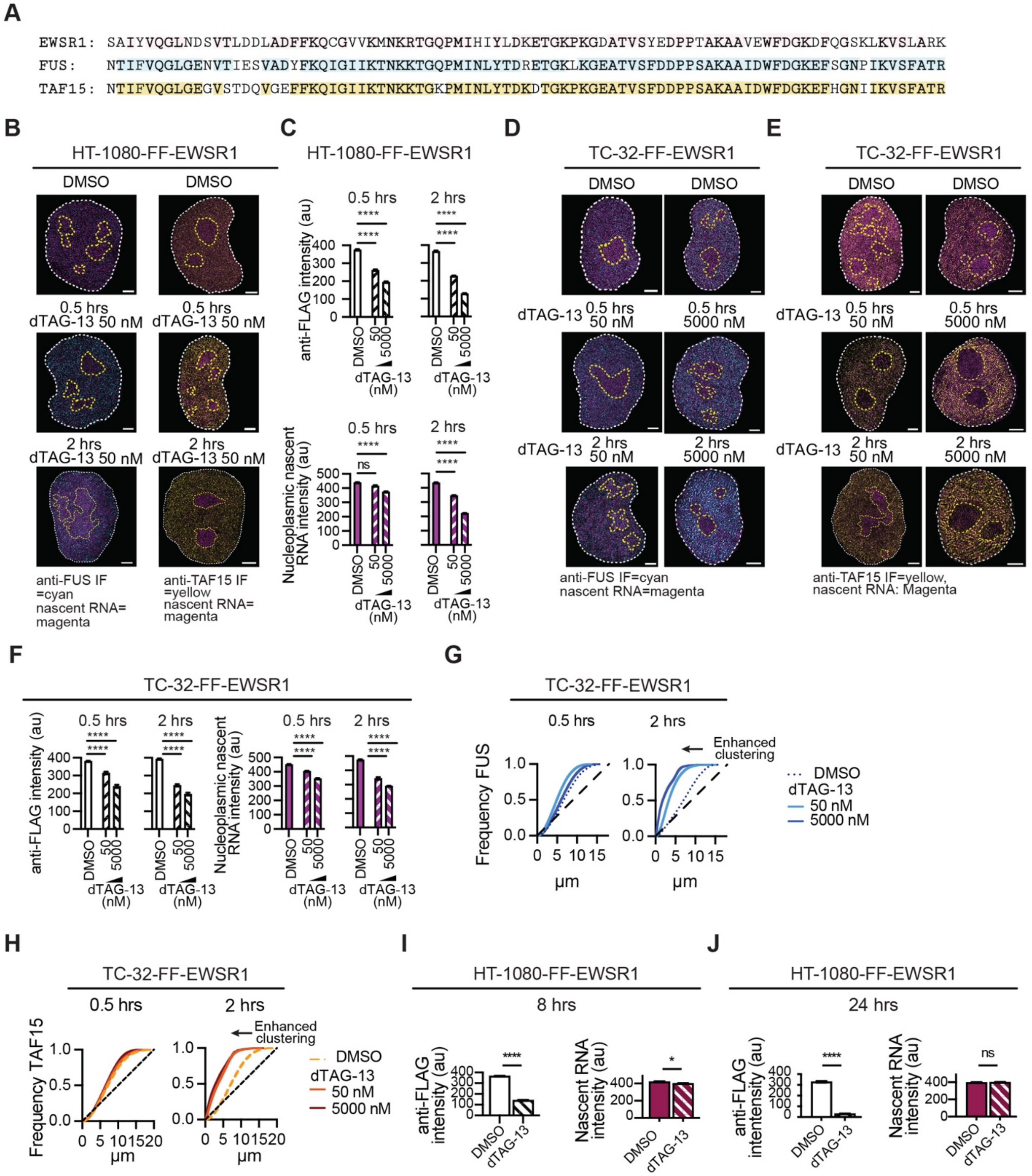


**Figure S9: FUS and TAF15 reorganize following EWSR1 degradation**

(**A**) The amino acid sequences of the FET protein RRM domains (extracted from UniProt).

(**B**) STED microscopy image of nuclei from HT-1080-FF-EWSR1 cells treated with DMSO or dTAG-13 (50 nM) for 0.5 or 2 hrs showing anti-FUS (cyan) (left panels) or anti-TAF15 (yellow)( right panels) Scale bar, 2 μm.

(**C**) Quantification of anti-FLAG IF intensity (upper) and nucleoplasmic nascent RNA intensity (lower) in HT-1080-FF-EWSR1 cells treated with DMSO or dTAG-13 (50 or 5000 nM) for 0.5 or 2 hrs.

(**D, E**) STED microscopy images of nuclei from TC-32-FF-EWSR1 cells treated with DMSO or dTAG-13 (50 or 5000 nM) for 0.5 or 2 hrs showing anti-FUS (cyan)(**D**) and anti-TAF15 (yellow)(**E**). Scale bar, 2 μm.

(**F**) Quantification of anti-FLAG IF intensity (left) and nucleoplasmic nascent RNA intensity (right) in TC-32-FF-EWSR1 cells treated with DMSO or dTAG-13 (50 or 5000 nM) for 0.5 or 2 hrs.

(**G, H**) Spatial cluster analysis of FUS (**G**) and TAF15 (**H**) pattern distributions. Frequency plots show analysis of STED microscopy images of a representative nucleus from TC-32-FF-EWSR1 cells treated with DMSO or dTAG-13 (5000 nM) for 0.5 or 2 hrs. The dashed line indicates a random point pattern distribution and arrowed solid lines indicate deviations a random point pattern distribution.

(**I, J**) Quantification of anti-FLAG IF intensity (left) and nucleoplasmic nascent RNA intensity (right) in HT-1080-FF-EWSR1 cells treated with DMSO or dTAG-13 (5000 nM) for 8 hrs (**I**) or 24 hrs (**J**).

(**A**, **D, E**) Images are representative of >20 nuclei per indicated FF-EWSR1 reporter cell line.

(**C, F, I, J**) Data are presented as mean ± SEM of 20 (**C,F**) or 15 (**I,J**) nuclei per treatment. Statistical significance was determined using one-way ANOVA * $p<0.05$, **** $p<0.0001$, ns non-significant.

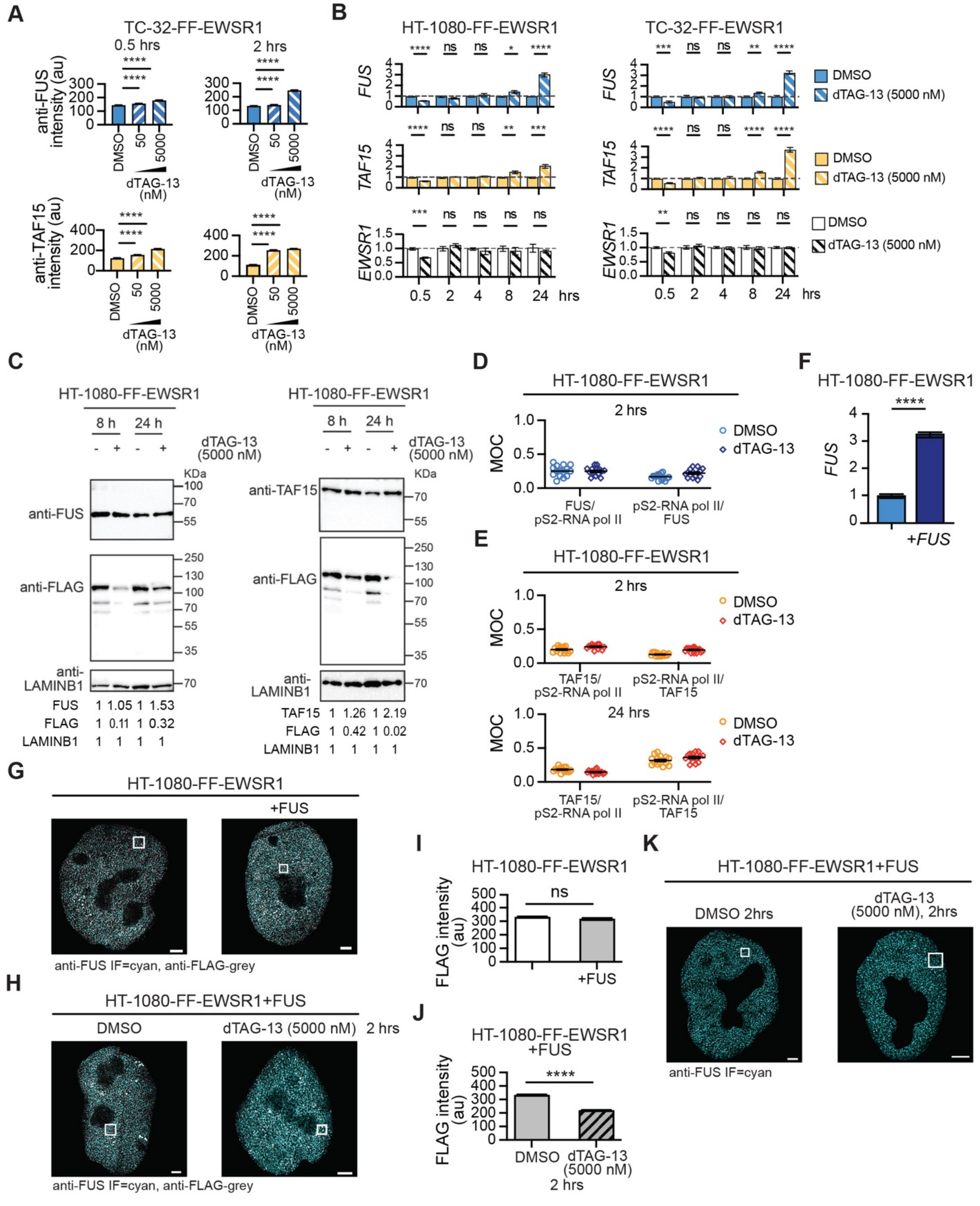


**Figure S10: The nuclear reorganization of FUS requires loss of EWSR1**

(**A**) Quantification of anti-FUS IF intensity (upper) and anti-TAF15 IF intensity (lower) in TC-32-FF-EWSR1 cells treated with DMSO or dTAG-13 (as indicated) for the times indicated.

(**B**) The expression of the indicated genes (*RPL27* normalized) following the addition of dTAG-13 (5000 nM) to HT-1080-FF-EWSR1 (left) or TC-32-FF-EWSR1 (right) cells at the indicated times. Data is shown relative to DMSO-treated cells.

(**C**) Immunoblot analysis of nuclear fractions lysates prepared from HT-1080-FF- EWSR1 reporter cell line treated with either DMSO or dTAG-13 (5000 nM) for 8 or 24 hrs and probed using the antibodies against the indicated proteins.

(**D**) Quantification of FUS colocalization with pS2-RNA pol II in HT-1080-FF-EWSR1 cells by MOC.

(**E**) Quantification of TAF15 colocalization with pS2-RNA pol II in TC-32-FF-EWSR1 cells by MOC.

(**F**) The expression of *FUS* (*RPL27* normalized) in HT-1080-FF-EWSR1 cells and HT-1080-FF-EWSR1 cells stably expressing *FUS*.

(**G, H**) STED microscopy images single channel (anti-FUS, cyan) and merged (anti-FUS, cyan; anti-FLAG, grey) of nuclei from HT-1080-FF-EWSR1 and HT-1080-FF-EWSR1+FUS cells (**G**) and HT-1080-FF-EWSR1+FUS cells treated with DMSO or dTAG-13 (5000 nM) (**H**).

(**I, J**) Quantification of anti-FLAG IF intensity in HT-1080-FF-EWSR1 and HT-1080-FF-EWSR1+FUS cells (**I**) and HT-1080-FF-EWSR1+FUS cells treated with DMSO or dTAG-13 (5000 nM) (**J**).

(**K**) STED microscopy images (anti-FUS IF (cyan)) of nuclei from HT-1080-FF-EWSR1+FUS cells treated with either DMSO or dTAG-13 (5000 nM) for 2 hrs.

(**A**) Data are presented as mean ± SEM of 20 nuclei per treatment.

(**B**) Data are presented as mean ± SEM of 6 replicates per treatment per time point. (**C**) Data are presented as mean ± SEM of 3 replicates. (**I, J**) Data are presented as mean ± SEM of 15 nuclei per treatment. Statistical significance was determined using one-way ANOVA. ⋆ p<0.05, ⋆⋆ p<0.01, ⋆⋆⋆ p<0.001, ⋆⋆⋆⋆ p<0.0001, ns non-significant.

(**G, H, K**) Images are representative of >15 nuclei per HT-1080-FF-EWSR1 or HT-1080 FF-EWSR1+FUS reporter cell line.
