## Supplementary material for "FUS and TAF15 safeguard the critical functions of the ribonucleoprotein network formed by EWSR1 and newly synthesized RNA": Resources and Methods

**KEY RESOURCES TABLE**

| REAGENT or RESOURCE | SOURCE | IDENTIFIER |
|---|---|---|
| **Antibodies** | | |
| Anti-EWSR1<br>[rabbit polyclonal, C-terminal region] | Thermo Fisher Scientific | Cat# PA5-35366;<br>RRID: AB_2552676 |
| Anti-FLI1<br>[rabbit polyclonal N-terminal region] | Abcam | Cat# ab15289;<br>RRID:AB_301825 |
| Anti-ERG<br>[rabbit monoclonal] | Abcam | Cat# ab110639;<br>RRID: AB_10864794 |
| Anti-ß-Actin<br>[rabbit polyclonal] | Cell Signaling Technology | Cat# 4967<br>RRID: AB_330288 |
| Anti-FLAG M2<br>[mouse monoclonal] | Sigma-Aldrich | Cat# F1804;<br>RRID: AB_262044 |
| Anti-pS2-RNA Pol II<br>[Anti-RNA polymerase II CTD YSPTSPS (phospho-S2) rabbit monoclonal] | Abcam | Cat# ab193468;<br>RRID:AB_2905557 |
| Anti-Vinculin<br>[mouse monoclonal] | Santa Cruz Biotechnology | Cat# sc-25336;<br>RRID:AB_628438 |
| Anti-FUS<br>[rabbit polyclonal] | ProteinTech | Cat# 11570-1-AP;<br>RRID: AB_2247082 |
| Anti-TAF15<br>[rabbit monoclonal] | Abcam | Cat# ab134916;<br>RRID: AB_2614922 |
| Anti-LaminB1<br>[rabbit polyclonal] | Thermo Fisher Scientific | PA5-19468<br>RRID: AB_10985414 |
| Anti-mouse IgG, HRP-linked Antibody | Cell Signaling Technology | Cat# 7076;<br>RRID: AB_330924 |
| Anti –rabbit IgG, HRP-linked antibody | Cell Signaling Technology | Cat# 7074;<br>RRID: AB_2099233 |
| Anti-rabbit IgG (H+L), F(ab')2 Fragment<br>[Alexa Fluor® 594 Conjugate] | Cell Signaling Technology | Cat# 8889;<br>RRID:AB_2716249 |
| Anti-mouse IgG (H+L), F(ab')2 Fragment<br>[Alexa Fluor® 594 Conjugate] | Cell Signaling Technology | Cat# 8890;<br>RRID: AB_2714182 |
| STAR RED, goat anti-rabbit IgG | Abberior | Cat# STRED-1002;<br>RRID:AB_2833015 |
| STAR RED, goat anti-mouse IgG | Abberior | Cat# STRED-1001;<br>RRID:AB_3068620 |
| **Chemicals and other reagents** | | |
| dTAG-13 | Tocris Biosciences | Cat# 6605,<br>CAS No: 2064175-41-1 |
| DMSO | Thermo Fisher Scientific | Cat# D12345 |
| DMEM | Thermo Fisher Scientific | Cat#11995073 |
| RPMI | Thermo Fisher Scientific | Cat# 11875093 |
| EMEM | ATCC | Cat# 30-2003 |
| Opti-MEM™ I Reduced-Serum Medium | Thermo Fisher Scientific | Cat# 31985070 |
| Trypsin-EDTA (0.25%), phenol red | Thermo Fisher Scientific | Cat# 25200056 |
| Phosphate Buffer Saline | Thermo Fisher Scientific | Cat# 10010023 |
| Fetal Bovine Serum (FBS) (Heat inactivated) | Thermo Fisher Scientific | Cat# 16140071 |
| Plasmocin Prophylactic | InvivoGen | Cat# Ant-mpp |

| | | |
|---|---|---|
| Nucleofector Kit T | Lonza | Cat# VCA-1002 |
| Nucleofector Kit R | Lonza | Cat# VCA-1001 |
| Novex™ Tris-Glycine Mini Protein Gels, 4–20%, 1.0 mm, WedgeWell™ format (10-well) | Thermo Fisher Scientific | Cat# XP04200BOX |
| Novex™ Tris-Glycine Mini Protein Gels, 4–20%, 1.0 mm, WedgeWell™ format (12-well) | Thermo Fisher Scientific | Cat# XP04202BOX |
| Novex™ Tris-Glycine SDS Running Buffer (10X) | Thermo Fisher Scientific | Cat# LC2675 |
| NuPAGE™ Tris-Acetate Mini Protein Gels, 3 to 8%, 1.0 mm, WedgeWell™ format, 12-well, 10 Gels/Box | Thermo Fisher Scientific | Cat# TA38012BOX |
| NuPAGE™ Tris-Acetate SDS Running Buffer (20X) | Thermo Fisher Scientific | Cat# LA0041 |
| iBlot™ 2 Transfer Stacks, PVDF, mini | Thermo Fisher Scientific | Cat# IB24002 |
| 4X Laemmli Sample Buffer | Bio-Rad | Cat# 1610747 |
| Nonfat dry milk | Cell Signaling Technologies | Cat# 9999 |
| Halt™ Protease and Phosphatase Inhibitor Cocktail (100X) | Thermo Fisher Scientific | Cat# 78440 |
| RIPA Lysis and Extraction Buffer | Thermo Fisher Scientific | Cat# 89901 |
| Pierce™ BCA Protein Assay Kits | Thermo Fisher Scientific | Cat# 23227 |
| RNAiMax | Thermo Fisher Scientific | Cat# 13778150 |
| Lipofectamine 3000 | Thermo Fisher Scientific | Cat# L3000015 |
| Trackit 100bp DNA ladder | Thermo Fisher Scientific | Cat# 10488058 |
| Ultrapure Agarose | Thermo Fisher Scientific | Cat# 16500500 |
| SybrSafe DNA gel stain | Thermo Fisher Scientific | Cat# S33102 |
| SuperSignal West Pico PLUS chemiluminescent substrate | Thermo Fisher Scientific | Cat# 34580 |
| Q5 HotStart 2X Mastermix | New England Biolabs | Cat# M0494 |
| GeneRuler 1kb plus DNA ladder | Thermo Fisher Scientific | Cat# SM1331 |
| DAPI | Thermo Fisher Scientific | Cat# 62248 |
| Ultrapure Glycerol | Thermo Fisher Scientific | Cat#15514011 |
| Puromycin dihydrochloride | Thermo Fisher Scientific | Cat# A1113803 |
| Blasticidin S HCl | Thermo Fisher Scientific | Cat# A1113903 |
| Geneticin selective antibiotic (G418 sulfate) | Thermo Fisher Scientific | Cat# 10131035 |
| Recovery™ Cell Culture Freezing Medium | Thermo Fisher Scientific | Cat# 12648010 |
| Ampicillin Ready-Made Solution | Quality Biological | Cat# 351-344-731 |
| Kanamycin Ready-Made Solution | Quality Biological | Cat# 120-349-731 |
| NheI -HF | New England Biolabs | Cat# R3131S |
| HindIII-HF | New England Biolabs | Cat# R3104S |
| One Shot™ TOP10 Chemically Competent E. coli | Thermo Fisher Scientific | Cat #C404010 |
| **Critical commercial assays** | | |
| MycoAlert Mycoplasma Kit | Lonza | Cat# LT07- 218 |
| AllPrep DNA/RNA Kit (QIAGEN) | Qiagen | Cat# 80204 |
| Maxwell RNA extraction kit | Promega | Cat# AS1390 |
| iScript Reverse Transcriptase cDNA synthesis kit | Bio-Rad | Cat# 1708840 |
| SiR-DNA staining kit | Cytoskeleton, Inc | Cat# CY-SC007 |
| Click-iT™ RNA Alexa Fluor™ 594 Imaging Kit | Thermo Fisher Scientific | Cat# C10330 |

| | | |
|---|---|---|
| NE-PER™ Nuclear and Cytoplasmic Extraction Reagents | Thermo Fisher Scientific | Cat# 78835 |
| CellTiter-Glo® Luminescent Cell Viability Assay | Promega | Cat# G7572 |
| **Deposited data** | | |
| Raw image files | This study | 10.17632/wk5tbr8j4f.1 |
| **Experimental models: Cell lines** | | |
| Human: HEK-293T | ATCC | Human Embryonic Kidney with SV-40 T antigen |
| Human: HT-1080 | ATCC | Fibrosarcoma |
| Human: A673 | ATCC | Ewing sarcoma *(EWSR1::FLI1*type1) |
| Human: TC-32 | Pediatric Oncology Branch, NIH | Ewing sarcoma *(EWSR1::FLI1*type1) |
| Human: SK-N-MC | Pediatric Oncology Branch, NIH | Ewing sarcoma *(EWSR1::FLI1*type1) |
| Human: TC-106 | Pediatric Oncology Branch, NIH | Ewing sarcoma *(EWSR1::ERG)* |
| **Oligonucleotides** | | |
| AllStars Neg. control siRNA (siNeg) | QIAGEN | Cat#1027281 |
| siEWSR1 (s4888)<br>5'-GCCUCCCACUGGUUAUACUtt-3' | Thermo Fisher Scientific Ambion Silencer Select | Cat# s4888 |
| siEWSR1 (s4886)<br>5'-AGAUUUUCAAGGGAGCAAAtt-3' | Thermo Fisher Scientific Ambion Silencer Select | Cat# s4886 |
| siFLI1 (s5266)<br>5'-CAAACGAUCAGUAAGAAUAtt-3' | Thermo Fisher Scientific Ambion Silencer Select | Cat# s5266 |
| siERG (s4812)<br>5'-CCACAGUGCCCAAAACUGAtt-3' | Thermo Fisher Scientific Ambion Silencer Select | Cat# s4812 |
| siFUS (s533595)<br>5'- AUUAUUAAGACAAAACACCGAtt-3' | Thermo Fisher Scientific Ambion Silencer Select | Cat# s533595 |
| siTAF15 (s15656)<br>5'-CCUAAUCCGUCAUGCGGAAtt-3' | Thermo Fisher Scientific Ambion Silencer Select | Cat# s15656 |
| *EWSR1*-Left HA primer new<br>5'-CCGGGGTTGCGAGATTTGCGC-3'<br>Template : gDNA | This study<br>**Fig. S1C** and **Fig. S2A** | |
| *EWSR1*-Right HA primer new<br>5'-GTTTGGGCGTTCCGGCTACCGCC-3' | This study<br>**Fig. S1C** and **Fig. S2A** | |
| Puromycin Forward<br>5'-GGACCGCGCACCTGGTGCATG-3'<br>Template: cDNA | This study<br>**Fig. S2B** | |
| *EWSR1*-Exon9-Reverse-mRNA<br>5'TGCCCATAAACACCCATGCTA-3'<br>Template: cDNA | This study<br>**Fig. S2B** | |
| *FLI1*-Exon 6-Reverse-mRNA<br>5'GTTATTGCCCCAAGCTCCTCT-3'<br>Template: cDNA | This study<br>**Fig. S2B** | |

| | | |
|---|---|---|
| *EWSR1* Forward qPCR<br>5'-GGGTATGGCACTGGTGCTTAT-3' | This study<br>**Fig. S10B** | |
| *EWSR1* Reverse qPCR<br>5'-CAGACTGAGCTGCATAGGAGG-3' | This study<br>**Fig. S10B** | |
| *FUS* Forward qPCR<br>5'-ACGGACACTTAGGCTATGG-3' | This study<br>**Fig. S7A, Fig. S10B** | |
| *FUS* Reverse qPCR<br>5'-TACCGTAACTTCCCGAGGTG-3' | This study<br>**Fig. S7A, Fig. S10B** | |
| *TAF15* Forward qPCR<br>5'-TGACCAGCAGTCAGGCTAG-3' | This study<br>**Fig. S7B, Fig. S10B** | |
| *TAF15* Reverse qPCR<br>5-CTCACATCAGACGGTCATC-3' | This study<br>**Fig. S7B, Fig. S10B** | |
| *RPL27* Forward qPCR<br>5'-GCAAGAAGAAGATCGCCAAG-3' | This study<br>**Fig. S7B, Fig. S10B** | |
| *RPL27* Reverse qPCR<br>5'- GACGACAGTTTTCTCCAAGG-3' | This study<br>**Fig. S7B, Fig. S10B** | |
| **Recombinant DNA** | | |
| Plasmid: pcDNA3.1(+)-*EWSR1* | This Study | |
| Plasmid: pReceiver-CMV-FUS-C-term Flag | GeneCopoeia | EX-F0952-M35 |
| **Software and algorithms** | | |
| Fiji (v1.52p) | Schindelin, J et al, doi:10.1038/nmeth.2019 | https://imagej.net/downloads |
| Spatial Statistics-Fiji Plugin | Andrey P et al, https://doi.org/10.1371/journal.pcbi.1000853<br>Ollion J et al, https://doi.org/10.1093/bioinformatics/btt276 | https://imagejdocu.list.lu/plugin/analysis/spatial_statistics_2d_3d/start |
| Coloc2-Fiji Plugin | Schindelin, J et al, doi:10.1038/nmeth.2019 | https://imagej.net/plugins/coloc-2 |
| RStudio (2025.09.1 Build 401) | RStudio Team (2019). RStudio: Integrated Development for R. RStudio, Inc., Boston, MA | http://www.rstudio.com/. |
| Graphpad Prism | | https://www.graphpad.com/ |
| FlowJo | FlowJo™ Software Windows,Version 10. BD Biosciences, Ashland, OR | https://flowjo.com/ |
| BioRender | | https://www.biorender.com/ |

### EXPERIMENTAL MODEL DETAILS

#### Cell lines and culture conditions

A673 and HEK-293T cells (ATCC, Manassas, Virginia) were cultured in Dulbecco's Modified Eagle Medium (DMEM; Thermo Fisher Scientific, Waltham, MA). TC-32, SK-N-MC and TC-106 cells (gifts from the Pediatric Oncology Branch, CCR) were cultured in RPMI-1640 (Thermo Fisher Scientific). HT-1080 (ATCC) cells were cultured in Eagle's Minimum Essential Medium (EMEM; ATCC). All media were supplemented with 10% fetal bovine serum (FBS, Thermo Fisher Scientific) and Plasmocin Prophylactic (InvivoGen, San Deigo, CA). Cells were maintained at 37°C in a humidified incubator with 5% $CO_2$.

Cell line identities were confirmed by short tandem repeat (STR) analysis (ATCC), and cultures routinely monitored for mycoplasma contamination using the MycoAlert Plus system (Lonza, Walkersville, MD). Generation of knock-in reporter cell lines and ectopic expression of plasmids are described below in **Method Details**.

### METHOD DETAILS

#### Generation of EWSR1 reporter cell lines

CRISPR-Cas9-mediated knock-in of reporter constructs at the 5' end of *EWSR1*-exon 1 was performed using two single-guide RNAs (sgRNAs), IVT-1001 and IVT-1002. sgRNAs were expressed from individual plasmids: pCE0590 (Lenti-SpCas9-2A-mCherry-EWSR1-IVT-1001) and pCE0591 (Lenti-SpCas9-2A-mCherry-EWSR1-IVT-1002). Two donor plasmids were used: pCE0567 and pCE0585.

The pCE0567 donor plasmid encodes a flexible G4S3 linker (GGGGS×3) fused to the N terminus of the mNeonGreen (mNG) fluorescent reporter. The pCE0585 donor plasmid encodes a G4S3 linker followed by a 3×FLAG epitope, the FKBP12$^{F36V}$ degron, and a puromycin resistance cassette linked via a self-cleaving P2A peptide (ATNFSLLKQAGDVEENPGP). Both donor plasmids contained ~800 bp homology arms flanking the predicted Cas9 cut site, with silent mutations introduced within the right homology arm to prevent re-cutting (see **Fig. S1A**).

Plasmids were transfected by lipofection or electroporation. HEK-293T cells were transfected using Lipofectamine 3000 (Thermo Fisher Scientific) and 2 µg total DNA in Opti-MEM™ I Reduced-Serum Medium following manufacturer's instructions (Thermo Fisher Scientific). All other cell lines were electroporated using the Lonza Nucleofector system with the following parameters: HT-1080 ($1\times10^6$ cells, 2 µg DNA, Kit T, program L-005); A673 ($1\times10^6$ cells, 2 µg DNA, Kit R, program A-028); TC-32 ($0.5\times10^6$ cells, 2 µg DNA, Kit R, program D-032); SK-N-MC ($1.2\times10^6$ cells, 2 µg DNA, Kit R, program D-032); TC-106 ($1.2\times10^6$ cells, 2 µg DNA, Kit R, program D-032).

Successfully modified clones were enriched either by fluorescence-activated single-cell sorting based on mNeonGreen expression or by puromycin selection followed by single cell sorting at the following concentrations: 1 µg/mL for HEK-293T, HT-1080, and A673, and 2 µg/mL for TC-32 cells. Puromycin selection was discontinued once selected single-cell clones were fully validated.

All plasmids were generated and sequence-verified by the Genome Modification Core, Laboratory Animal Sciences Program, Frederick National Laboratory for Cancer Research, NCI, NIH.

#### Generation of EWSR1- and FUS-expressing FF-EWSR1 cell lines

For ectopic expression of EWSR1, the *EWSR1* CDS (ENST00000397938) was synthesized with flanking NheI and HindIII restriction sites (Genewiz, South Plainfield, NJ) and cloned into

NheI/HindIII-digested pcDNA3.1(+) (Thermo Fisher Scientific). Following bacterial transformation, plasmid DNA was isolated using the PureLink HiPure Miniprep Kit (Thermo Fisher Scientific), and full-length sequence integrity was confirmed by whole-plasmid Oxford Nanopore sequencing (CCR Genomics Core, CCR, NCI).

The pcDNA3.1(+)-EWSR1 plasmid DNA was transfected into HEK-293T and HT-1080-FF-EWSR1 cells using 2 µg DNA and Lipofectamine 3000 in Opti-MEM™ I Reduced-Serum Medium. Forty-eight hours (hrs) post-transfection, cells were selected using Geneticin (G418; 1 mg/mL, Thermo Fisher Scientific) until stable populations were obtained.

For ectopic expression of FUS, 2 µg plasmid DNA expressing the *FUS* cDNA from the CMV promoter (Genecopoeia (Rockville, MD) was transfected into HT-1080-FF-EWSR1 cells using Lipofectamine 3000 in Opti-MEM™ I Reduced-Serum Medium following manufacturer's instructions (Thermo Fisher Scientific). Forty-eight hrs post-transfection, stable cell populations were generated by selection with G418 (1 mg/mL).

**Sequence analysis of modified cells**

Genomic DNA and total RNA were isolated from genetically modified clones and parental controls using the QIAquick AllPrep DNA/RNA Kit (Qiagen, Germantown, MD) according to the manufacturer's instructions.

Knock-in events were validated by PCR amplification of genomic DNA using primers spanning the targeted integration site and Q5 Hot Start 2× Master Mix (New England Biolabs (NEB) Ipswich, MA). PCR products were resolved on 6% TBE polyacrylamide gels, stained with SYBR Safe DNA dye, and visualized using the Omega Lum G imaging system (Aplegen, Pleasanton, CA). Amplicons were sequenced using primers located external to both homology arms to confirm precise genomic insertion.

Allelic insertion of the puromycin–FKBP12$^{F36V}$–FLAG cassette was assessed using transcript-specific PCR. First-strand cDNA was synthesized from 1 µg total RNA using the iScript cDNA Synthesis Kit (Bio-Rad, Hercules, CA), and PCR amplification used a forward primer mapping to the puromycin cassette and reverse primers specific to either *EWSR1*-exon 9 or *FLI1*-exon 6, and Q5 Hot Start 2× Master Mix. PCR products were resolved and visualized as described above.

**RNAi interference**

siRNA-mediated gene silencing was performed using RNAiMax (Thermo Fisher Scientific). Cells were transfected at a final siRNA concentration of 20 nM. HEK-293T and EWS cells were transfected at $1.35\times10^5$ cells per well of a 6-well plate were transfected using 4.5 µL RNAiMAX. HT-1080 cells were transfected at $2\times10^5$ cells per well using 4 µL RNAiMax. Analyses were performed 48 hrs post-transfection.

**Flow cytometry**

Cells were washed with ice-cold PBS and resuspended in FACS buffer (phenol red–free medium with 2% FBS). DAPI (1:1000) was used for live/dead discrimination. Samples were filtered through a 35 µm mesh and analyzed on a Sony ID7000 spectral analyzer, collecting ≥8000 events per sample. Data were analyzed using FlowJo. Sequential gating was applied to select singlets and live cells, followed by mNeonGreen fluorescence quantification. Median fluorescence intensity was recorded directly from the ID7000 output.

**Immunoblotting**

Whole-cell lysates were prepared in radioimmunoprecipitation (RIPA) Lysis and Extraction Buffer (Thermo Fisher Scientific) supplemented with 1x Protease and Phosphatase Inhibitor Cocktail (Thermo Fisher Scientific) followed by sonication for 10-20 secs and clarification by centrifugation at 10000 rpm for 10 minutes (min) at 4°C. Protein concentrations were determined using the Pierce™ BCA Protein Assay Kit (Thermo Fisher Scientific). Equal amounts of protein (25–30 μg) were resolved on precast 4 to 20% SDS–PAGE or 3-8% Tris-Acetate gels and transferred to PVDF membranes using the iBlot2 (Thermo Fisher Scientific) semi-dry transfer device.

For nuclear fractionation, cells treated with DMSO or dTAG-13 for the indicated time points were harvested and pelleted. Nuclear and cytoplasmic extracts were prepared using the NE-PER™ Nuclear and Cytoplasmic Extraction Kit (Thermo Fisher Scientific) according to the manufacturer's instructions, with the inclusion of an additional wash of the nuclear pellet to minimize cytoplasmic contamination. Nuclear protein concentrations were determined by BCA assay, and 20 μg of nuclear protein was resolved and transferred to PVDF membranes as described above.

Membranes were blocked in 5% non-fat dry milk in TBS-T for one hour at RT and incubated with primary antibodies overnight at 4°C. The following antibody dilutions were used: anti-EWSR1 (1:1000), anti-FLI1 (C-terminal; 1:1000), anti-ERG (1:500), anti-Vinculin (1:1000), anti-Lamin B1 (1:5000), anti-FLAG M2 (1:3000), anti–phospho-Ser2 RNA polymerase II (1:3000), anti-FUS (1:2000), anti-TAF15 (1:1000), and anti-β-Actin (1:1000). After washing thrice with TBS-T, membranes were incubated with HRP-conjugated secondary antibodies and developed using SuperSignal West Pico PLUS chemiluminescent substrate (Thermo Fisher Scientific). Signals were visualized using the Omega Lum C imaging system.

**dTAG-13 treatment**

A 1000× stock of dTAG-13 was prepared in DMSO. Cells were treated with either vehicle (0.001% DMSO) or dTAG-13 at the indicated concentrations for the indicated durations prior to analysis.

**Gene expression analysis**

Total RNA was extracted using the Maxwell SimplyRNA system (Promega). cDNA synthesis and qPCR were performed as described above. Relative expression was calculated using the ΔΔCt method, with gene-specific expression normalized to Ribosomal Protein L27 (*RPL27*).

**ATP metabolic activity assay**

Cells were transfected in white 96-well plates (1,800 cells/well) with siRNAs at a final concentration of 20 nM. Forty-eight hours post-transfection, cells were treated with DMSO or dTAG-13 at the indicated concentrations. Cell viability was assessed using CellTiter-Glo (Promega), and luminescence was measured on an EnSight multimode plate reader (PerkinElmer). Raw luminescence values were normalized to siNeg/DMSO-treated controls and plotted as mean ± standard error of mean (SEM) of six replicate transfections.

**Immunofluorescence sample preparation**

For IF experiments, $1×10^5$ cells were seeded per well of a 6-well plate on round, sterile glass coverslips and allowed to adhere overnight. Cells (untreated or treated) were processed for imaging as follows. Culture medium was aspirated, and coverslips were washed once with PBS, followed by fixation with 4% paraformaldehyde for 3 min at room temperature. After fixation, coverslips were washed three times with PBS and permeabilized with 0.5% Triton X-100 for 3 min at room temperature. Coverslips were then washed three additional times with PBS and blocked for 1 hour (hr) at room temperature in 10% normal goat serum (Thermo Fisher Scientific). Primary antibodies were diluted in blocking buffer and incubated with coverslips overnight at 4°C. Coverslips were subsequently washed three times with PBS and incubated with the appropriate secondary antibodies for 4 hrs at room temperature in the dark. Following secondary antibody incubation, coverslips were mounted using mounting medium containing 90% glycerol, 100 mM Tris (pH 8.0), and 0.1 mg/mL *p*-phenylenediamine. For STED-based imaging, mounting medium lacking antifade (*p*-phenylenediamine) was used. Coverslips were sealed with clear nail polish and cured overnight in the dark prior to imaging. Parental, unmodified cells were used as background controls for the mNeonGreen (488 nm) channel, and secondary antibody–only controls were used to assess background signal for all other fluorescence channels. Across all IF experiments, imaging parameters were held constant, and laser power and detector settings were optimized to prevent signal saturation.

**DAPI, SiR-DNA and nascent RNA visualization**

For DAPI staining, all steps up to and including secondary antibody incubation were performed as described above. Following secondary antibody washes with PBS and prior to mounting, coverslips were incubated with DAPI (Thermo Fisher Scientific; 1:1000 dilution in PBS) for 3 min at room temperature. Coverslips were then washed three times with PBS, mounted, and cured in the dark prior to imaging.

For SiR-DNA labeling, $1 \times 10^5$ cells per well were seeded on glass coverslips in 6-well plates and allowed to adhere overnight. SiR-DNA (Spirochrome, Thurgau, Switzerland) was added to the culture medium at a final concentration of 500 nM in 2 mL of complete medium, together with 5 µM verapamil. Cells were incubated for 1 hr at 37°C before proceeding with fixation and sample preparation as described above.

For nascent RNA visualization, $1 \times 10^5$ cells per well were plated on coverslips in 6-well plates and allowed to attach overnight. Cells were incubated with 5-ethynyl uridine (EU) at a final concentration of 1 mM in 2 mL of culture medium for 40 min at 37°C. Following incubation, cells were processed according to the manufacturer's instructions using the Click-iT™ RNA Alexa Fluor™ 594 Imaging Kit (Thermo Fisher Scientific).

**Confocal microscopy**

Cells were plated and prepared as described above. For confocal imaging, post incubation with secondary antibody, coverslips were further incubated with DAPI (Thermo Fisher Scientific) for 3 min in the dark before mounting onto clean glass slides. Slides were imaged in the confocal mode, on a Nikon SoRa spinning disk microscope equipped with a photometrics BSI sCMOS camera with the 20X objective lens. At least 20 nuclear z-stacks of step size 0.2 µm were obtained per image covering a nuclear volume of ~4 µm in each channel. Images, obtained in the .nd2 format, were imported into Fiji (v1.52) and a representative field of view of merged signals was generated.

**Fluorescence Recovery After Photobleaching (FRAP)**

For fluorescence recovery after photobleaching (FRAP) analysis, $5 \times 10^4$ cells were seeded in 35-mm glass-bottom dishes (Ibidi USA Inc., Fitchburg, WI) and allowed to adhere overnight. Prior to imaging, culture medium was replaced with phenol red–free medium supplemented with 10% fetal bovine serum. For each nucleus analyzed, three regions were defined: a 3 μm region outside the nucleus to measure background fluorescence, a region within the nucleus containing fluorescent signal to serve as a reference, and a 3-μm region within the nucleus designated as the region of interest (ROI). Photobleaching was performed using the 488-nm laser line set to 80% laser power. Following bleaching, images were acquired every 5 s for a total duration of 5 min using a Zeiss LSM 880 Airyscan confocal microscope equipped with a GaAsP spectral detector and an argon laser at the indicated wavelength. Background-subtracted fluorescence intensities for the reference and ROI regions were extracted using Zeiss ZEN (black) software. ROI intensities were normalized to the reference signal using the FRAP module within ZEN (black). Recovery curves were fit using a double-exponential model, and dissociation constants were obtained as direct outputs of the software. All analyses and curve fitting were performed using the FRAP module in ZEN (black) with default settings. Normalized recovery curves and dissociation constants were exported to GraphPad Prism, where FRAP curves are presented as mean values and dissociation constants are reported as mean ± SEM.

**Super resolution confocal microscopy (Spinning Disk SoRa)**

Slides prepared as described above were imaged on a Nikon SoRa spinning disk microscope equipped with a photometrics BSI sCMOS camera with the 60X Apo TiRF objective lens (Numerical Aperture: 1.49) in the SoRa mode. Z-stacks across the nuclear volume (~5 μm) was collected with a step size of 0.2 μm. Post image collection, batch processing was performed using standard macros in the NIS software for channel alignment and deconvolution. Pre-processed images were imported into Fiji to build single channel and merged images of representative nuclei.

For line plots, single channel SoRa images from the indicated representative insets were obtained from the merged images. The inbuilt “Plot profile” function in Fiji was used to generate the line profile across the marked distance in the inset with no changes to any parameters within the function. To determine nuclear intensity, DAPI was used to generate a nuclear mask, and integrated density was calculated on the masked regions using standard functions in Fiji for each channel. Average integrated density with SEM was plotted as shown.

**Stimulated Emission Depletion Microscopy (STED)**

Coverslips and slides were prepared as described above and mounted using antifade-free mounting medium. STED imaging was performed on a Leica Stellaris 5 STED microscope equipped with four HyD S detectors, one HyD X STED detector, and a 775-nm STED depletion laser. Images were acquired using a HC plan-apochromat STED White 100x (N.A. 1.4) oil immersion objective lens in confocal, TauSTED, and TauSTED-Xtend acquisition modes. Central z-planes encompassing whole nuclei were imaged at a final lateral resolution of ~50 nm, and higher-magnification insets were acquired at a final resolution of ~20 nm.

Acquired images were further deconvolved using the Lightning processing algorithm (standard function) in the Leica NIS software. Secondary antibody–only controls were used to perform background subtraction for each relevant fluorescence channel. Pre-processed images (.lif files) were imported into Fiji for generation of representative single-channel and merged images.

For quantification of total nuclear signal from STED images, background-subtracted images were imported into Fiji, and the integrated density function was used to calculate total fluorescence intensity for each channel of interest across individual nuclei. Quantitative data were exported to GraphPad Prism and are presented as mean ± SEM for the indicated number of replicates.

To define low- and high-intensity FF-EWSR1 signal populations, background-subtracted STED images were analyzed in Fiji. For each image, intensity thresholds were set at fivefold above the minimum signal above background. A size exclusion threshold of 20 pixels was applied using the "Analyze Particles" function to identify discrete foci. The number of foci and their linear dimensions were quantified using the "Measure Particles" function. Resulting measurements were exported to GraphPad Prism and plotted as mean ± SEM for the indicated replicates. All analyses were performed using default software and function settings in Fiji unless otherwise specified.

### STATISTICAL ANALYSIS

#### Ripley's *k*-statistic

Deconvolved and background subtracted STED images were used to obtain either frequency or cumulative frequency curves. The spatial statistics plugin was downloaded from https://mcib3d.frama.io/3d-suite-imagej/ and installed into Fiji. Single channel frequency curves were built using binarized (using the inbuilt "make binary" function) 8-bit images as input and using the simple statistics option within the plugin. All default settings were used.

To obtain cumulative frequency curves, single channel 8-bit images were generated from the two channels of interest. The nucleic acid/protein signal to be used as a reference structure was first converted to a mask (using the "convert to mask" inbuilt function) and the other channel was binarized as described above. Spatial statistics option was used, and the appropriate channel was used for the reference structure.

All frequency curves were obtained as the H-transformation of the Ripley's *k*-statistic. The output from the Fiji plugin was exported as a .csv file. In the case of protein signals, outputs exceeded $1 \times 10^6$ measurements per nucleus, and in the case of nucleic acids, over $1 \times 10^8$ measurements per nucleus. To plot frequency or cumulative frequency curves, every 999th data point from the .csv data file was extracted using RStudio and the code below. The subset of the data was then plotted in Graphpad prism to generate the curves shown.

```
>setwd("C:/Users/ STED") # set working directory where data is present
>data=read.csv("datafile1.csv",header=TRUE,sep=",") #read the .csv file generated by the plugin in Fiji
>data_subset=data[seq(999, nrow(data), by = 999), ] #extract every 999th row from the .csv datafile
>head(data_subset) # shows the first 6 rows of the newly subsetted data file
>write.csv(data_subset,"data_subset.csv",row.names=FALSE)#rewrite the subset data file as a .csv to import into Graphpad Prism.
```

In all the frequency and cumulative frequency plots the dashed line represents a random point pattern distribution. In each plot, arrowed solid lines indicate deviations from a random point pattern distribution and arrowed dotted lines indicate a random distribution.

#### Pearson correlation coefficient (PCC) and Mander's overlap coefficient (MOC)

Deconvolved and background-subtracted SoRa or STED images were used for the calculation of these colocalization metrics. For 2D Pearson correlation coefficient (PCC) and Mander's overlap coefficient (MOC) analyses of different channels, the "Coloc2" plugin in FIJI (v1.52) was downloaded from https://github.com/fiji/Colocalisation_Analysis and utilized. Briefly, the central plane of each nuclear image was used and single channels were split. Each of the relevant single channel images was used as input for the “Coloc2” plugin to obtain the required values with thresholding values based on secondary only controls for each channel. The data was exported to GraphPad Prism and plotted as depicted.

**Statistical analysis of significance**

Results shown as mean and standard error of the mean (SEM) were calculated using GraphPad Prism version 10.6.1, GraphPad Software, San Diego, California USA, www.graphpad.com. Statistical analysis was conducted using an unpaired t-test with Welch’s correction or One-way ANOVA in GraphPad Prism. A p-value of $<0.05$ was considered significant.
